# Spring-loaded inverted pendulum goes through two contraction-extension cycles during the single stance phase of walking

**DOI:** 10.1101/509687

**Authors:** Gabriel Antoniak, Tirthabir Biswas, Nelson Cortes, Siddhartha Sikdar, Chanwoo Chun, Vikas Bhandawat

## Abstract

Despite the overall complexity of legged locomotion, the motion of the center of mass (COM) itself is relatively simple, and can be qualitatively described by simple mechanical models. The spring-loaded inverted pendulum (SLIP) is one such model, and describes both the COM motion and the ground reaction forces (GRFs) during running. Similarly, walking can be modeled by two SLIP-like legs (double SLIP or DSLIP). However, DSLIP has many limitations and is unlikely to serve as a quantitative model for walking. As a first step to obtaining a quantitative model for walking, we explored the ability of SLIP to model the single stance phase of walking across the entire range of walking speeds. We show that SLIP can be employed to quantitatively model the single stance phase except for two exceptions: first, it predicts larger horizontal GRFs than empirically observed. A new model - angular and radial spring-loaded inverted pendulum (ARSLIP) can overcome this deficit. Second, even the single stance phase has active elements, and therefore a quantitative model of locomotion would require active elements. Surprisingly, the leg spring undergoes a contraction-extension-contraction-extension (CECE) during walking; this cycling is partly responsible for the M-shaped GRFs produced during walking. The CECE cycle also lengthens the stance duration allowing the COM to travel passively for a longer time, and decreases the velocity redirection between the beginning and end of a step. A combination of ARSLIP along with active mechanisms during transition from one step to the next is necessary to describe walking.

## 1 Introduction

Legged locomotion is complex; therefore, many approaches to legged locomotion focus on the motion of the animal’s center of mass (COM) rather than on the detailed dynamics of each joint [18]. Traditionally, walking and running were described using different mechanical systems: The stiff-legged inverted pendulum (IP) was used as a model for walking [22, 44, 10], while the spring-loaded inverted pendulum (SLIP) with its compliant legs was used as a model for running [6, 33, 8, 2, 15, 36, 39]. Recent realization that legs are compliant during walking as well [29,10] led to the development of the double SLIP (DSLIP) model, in which each leg of a biped is modeled as a spring. DSLIP extends SLIP with a double stance phase during which the COM is supported by two “springy” legs [19,38]. These studies suggest that simple mechanical models can serve as conceptual models for locomotion. However, a central question is whether these models describe locomotion well enough to serve not just as conceptual models, but as quantitative models for locomotion that can be used to understand the control of locomotion by the nervous system [24].

Current versions of DSLIP models have many deficiencies which make it unlikely that they can serve as a quantitative model for walking in their current form. As noted by the authors themselves, DSLIP finds stable gaits only for a limited range of speeds [19, 30], and predicts within-step variations in COM height and GRFs [30, 25] that are far greater than those observed during locomotion. DSLIP is also unable to reproduce experimentally observed stance durations; predicted stance durations are always shorter than those observed empirically. These deficiencies of DSLIP might arise from two sources. First, we have already shown that replacing each leg with a single linear spring is an oversimplification, one needs to also account for tangential forces [5]. Additionally, it is possible that the right parameter regime that models experimentally observed dynamics most accurately, particularly in the case of walking, have not been discovered. Thus, it is likely that either a more rigorous approach to parameter search, or a different dynamical model, or a combination of the two would yield a quantitative model for walking.

The dynamical model proposed here is the angular and radial spring loaded pendulum (ARSLIP) model (Fig. 1A), and is aimed at improving the ability of SLIP to model ground reaction forces (GRFs) experienced by animals during walking. Animals receive GRFs in the form of normal and frictional forces that act vertically and horizontally, respectively (Fig. 1B). SLIP assumes that these two forces adjust such that the total GRF is always aligned along the effective leg (radial forces). There is no *a priori* reason that GRFs should always be along the leg, and indeed several studies have indicated otherwise [31, 35]. ARSLIP extends SLIP by providing a mechanism for modeling tangential forces. The tangential forces are restorative and changes sign at mid-stance, see Fig. 1; these restorative forces were shown to be important for walking [5]. There are other models for tangential forces. But most of these models only produce unidirectional torques [40,3,42]. Some models whose intended purpose is different to ours do produce tangential forces similar to the ones produced in ARSLIP. One method adds a roller foot to a springy leg [45, 28]. Another model proposes that the COM is below the point through which the force acts, resulting in restoring forces [31, 41]. Both of these methods help in overcoming some limitations of SLIP, but because the radial and tangential forces are not independent, many weaknesses remain. In ARSLIP, the radial and tangential forces are independently tuned by two independent springs. Conceptually, a simple method to visualize the model is to think of an angular spring connecting the foot to the leg as depicted in Fig. 1A. More details about the model and its actualization can also be found in Supplementary materials (Section A).

**Figure 1:**
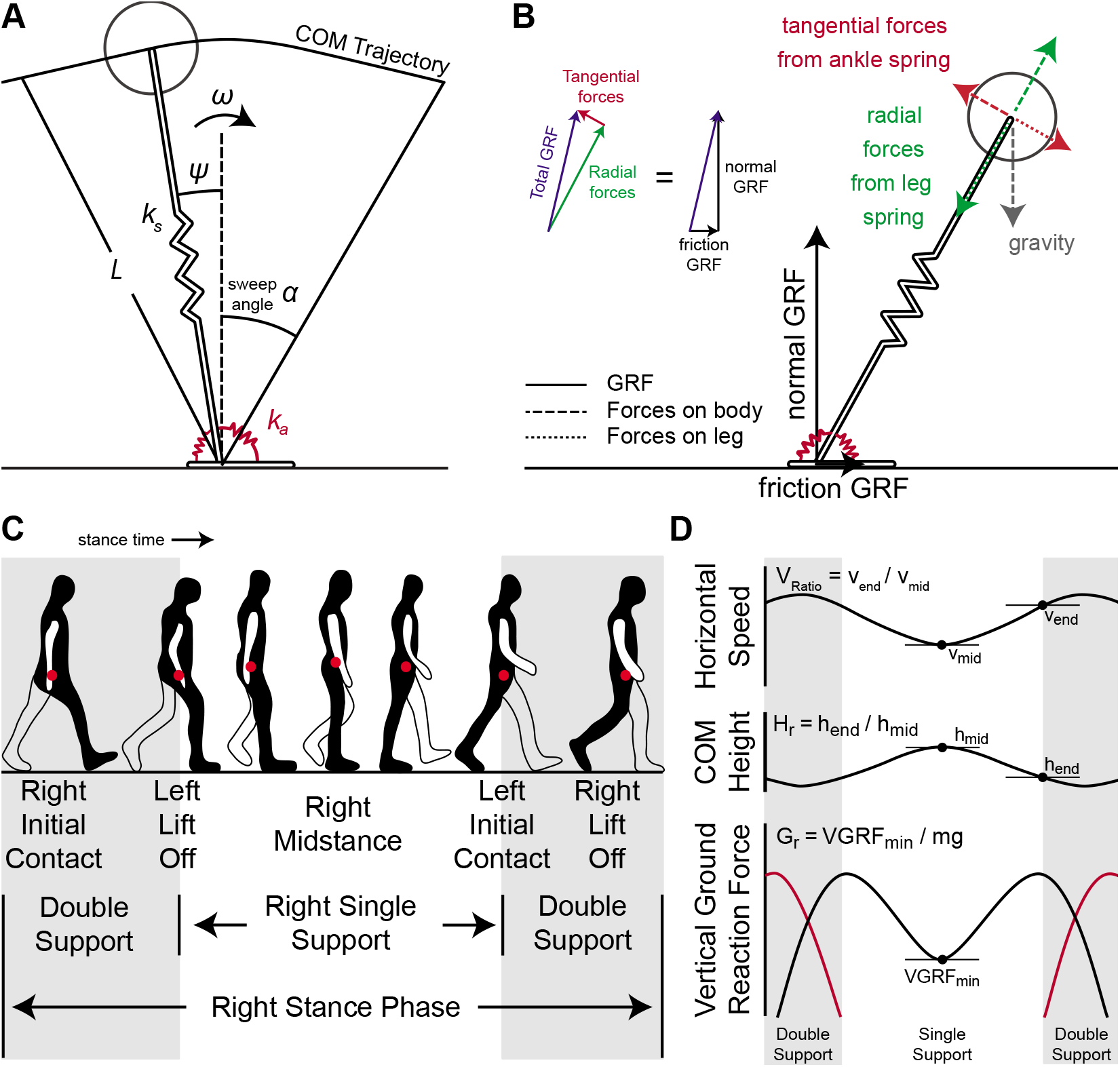
ARSLIP template and gaitspace constraint description. **A.** The ARSLIP template contains a leg spring of stiffness *k*_*s*_ and an ankle angular spring of stiffness *k*_*a*_ that resists motion away from the vertical orientation of the leg. **B.** The interaction between the ground, leg, and the body in the ARSLIP template. The GRFs that the leg recieves are denoted by the black arrows. The forces are then transmitted to the body by the leg in the form of the dashed tangential and radial forces, colored red and green respectively. The equal and opposite forces that the leg recieves from the body, the red and green dotted lines, cancel the GRFs as shown in the inset, since the leg is massless. **C.** Visualization of the stance period of gait for the right leg. The red dot represents the center of mass. Single support occurs in the middle of the stance phase as is when a single leg is in contact with the ground. **D.** Visualization of the constraints on the gaitspace. The velocity constraint is the ratio of the horizontal velocity at the end of single support to the midstance horizontal velocity. The height constraint is the ratio of the COM height at the end of single support to the midstance height of the COM. The vertical ground reaction force constraint is the minimum ground reaction force during single support normalized by body weight.

Our approach to evaluating a model is different from previous studies in two respects: The first point is conceptual. In contrast to most traditional approaches to simplified “template” models which are passive, we conceptualize our models as passive within a step. That is, the model parameters are the parameters that are controlled by an animal during a step.

The second point relates to defining what constitutes a successful model. Most studies focus on a single aspect of locomotion such as GRFs; our approach focuses on three important aspects of locomotion: GRFs, COM kinematics, and stance duration. That is, a successful model must produce realistic GRFs within the constraints of experimentally observed COM kinematics over experimentally observed stance duration. These three constraints are rarely satisfied [31, 30, 32] simultaneously in most studies of locomotion. Without satisfying the constraints on force, kinematics and duration simultaneously, the problem is under constrained: The COM height determines the natural timescale of the system (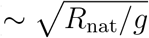); and, in understanding locomotion, one important consideration is how does the stance duration compare to the natural time constant. Allowing *R*_nat_ to be arbitrary ignores this important constraint. It is precisely because of the presence of the gravitational time scale that slow walking is a challenge that therefore has to be addressed in any realistic model of locomotion [5].

This manuscript focuses on the single support phase of walking Fig. 1C. As mentioned earlier, walking can be divided into two phases: a relatively short double support phase during which both legs are on the ground, and the longer single support phase. Neither the stiffness nor the natural length of the leg remains constant during the entire stance duration. During the double stance phase, the ankle is flexible; as the leg proceeds to its single stance phase, the ankle becomes stiffer. Moreover, a very influential series of studies have described the double support phase as a transition phase [16, 1] that is unlikely to be modeled using pendular models. Therefore, as a first step to obtaining a biomechanical model of the entire walking cycle, here we study the mechanics of the single support phase. Using non-dimensional analysis, we show that SLIP can model the single support phase of walking, and ARSLIP expands the parameter range over which realistic walking can be modeled. Surprisingly, fitting SLIP to empirical walking data shows that the stance leg goes through a contraction-expansion-contraction-expansion (CECE) cycle during single stance. This cycling appears important for reducing the extent to which velocity vectors have to be redirected during the double stance phase.

## 2 Methods and Results

### 2.1 Non-dimensional analysis of SLIP and ARSLIP

Successful models of locomotion have to be constrained not only by the average COM kinematics, but also by the within-step variations in the COM kinematics. For example, the small changes in COM height that is observed during a step (Fig.1D) places stringent constraints on the stiffness of the spring that can be employed. In this section, we will define constraints on the COM height, speed and GRFs (Fig.1D) that must be satisfied by the model. The parameter subspace within which these constraints are satisfied is called *gaitspace*. This analysis extends a previous analysis of SLIP [5] by including a new model, ARSLIP, and by including GRF in our analysis.

#### 2.1.1 Constraints on COM height and speed, as well as GRF

The COM height for a human during walking changes by less than 10% of its leg length [29]. We incorporate the constraints on height [5] by using the “height ratio”, *H*_*r*_ ≡ (Beginning COM Height)/(Mid-stance COM height), see Fig. 1D. The constraint on height was imposed as follows:

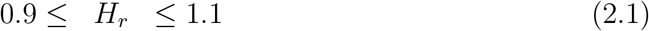

Changes in the horizontal speed of the COM lie within 25% of the mean [11, 7, 17]. Constraints on speed were incorporated by introducing a “velocity ratio”, *V*_*r*_ ≡ (Beginning horizontal velocity)/(Mid-stance horizontal velocity). The constraint on speed was imposed as follows:

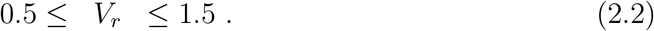

A relatively large band of speed is allowed because for some animals there is a large fluctuation in speed. Only symmetric walking gaits are considered so that the height and horizontal speed at the beginning and end of a given stance phase is the same.

We imposed two constraints on VGRF. First, VGRF rarely goes below 30% of the weight of the animal [25, 4]. To capture this constraint, we define “VGRF ratio”, *G*_*r*_ ≡ (minimum VGRF)/*mg*, and impose the constraint

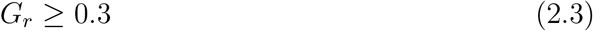

Second, for mammals, including humans [12, 29], certain birds [4, 14], and most quadrupeds [43, 9, 27], the VGRF has a mid-stance minimum and is flanked on each side by two local maxima thereby producing a characteristic M-shape, see Fig. 1C. We will evaluate how the models perform when we impose the condition that the VGRF, *F*_*y*_, has to have a minimum (“VGRF convexity”) at the midstance:

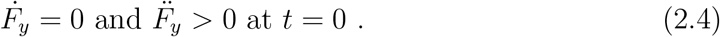

Realistic walking gaits must satisfy (2.1–2.4).

#### 2.1.2 Models, Evolution Equations and Dimensionless parameterizations

The SLIP and ARSLIP models are shown in Fig. 1. SLIP is merely the special case of ARSLIP with the angular-spring stiffness set to zero. The governing differential equations for single stance of ARSLIP can be obtained from energy considerations, see Supplement A, and are given by

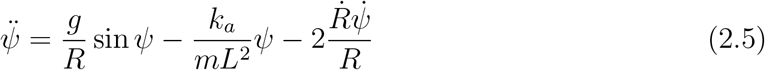

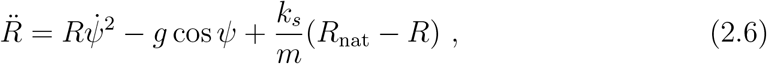

where *ψ*(*t*) is the angle the leg makes from the vertical mid-stance position, *L*(*t*) is the effective leg length from the COM to the contact point during mid-stance, *k*_*a*_ the stiffness of the angular spring, *k*_*s*_ the stiffness of the leg spring, *g* is the gravitational acceleration constant, and *R*_nat_ the uncompressed length of the effective leg. As compared to SLIP which only allows radially outward forces controlled by the spring constant, *k*_*s*_, the ARSLIP admits additional restoring tangential forces that are controlled by angular spring stiffness, *k*_*a*_. The evolution of the COM from the mid-stance position is assumed to be symmetric, so that the initial conditions for the differential equation are: *R*(0) = *R*_o_, *Ṙ*(0) = 0, *ψ*(0) = 0, and 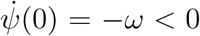; the mid-stance is chosen to occur at *t* = 0. Thus a symmetric gait is specified by six parameters: *k*_*s*_, *k*_*a*_, *R*_nat_, *R*_o_, *ω* and *α*, where *α* denotes the “sweep angle” (Fig. 1A).

To perform the analysis that can be generalized across different species we recast the evolution equation in terms of dimensionless variables:

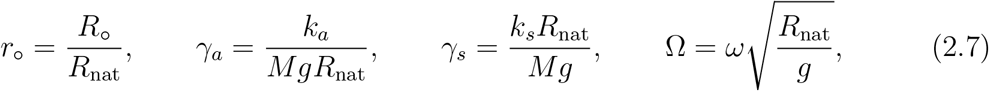

where *r*_o_ is the non-dimensional vertical height of the COM at mid-stance, *γ*_*a*_, *γ*_*s*_ represent the non-dimensional angular and radial spring constants respectively, and Ω is the non-dimensional angular speed at mid-stance. The evolution of the COM in non-dimensional form reads

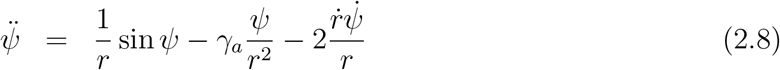

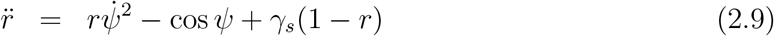

where 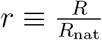 is the non-dimensional radial coordinate and the differentiations are now with respect to the non-dimensional time 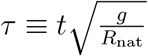. As is well known, working with dimensionless quantities also reduce the number of parameters that one needs to independently vary making it easier to explore the full parameter space. In terms of dimensionless variables, SLIP has four parameters, *γ*_*s*_, Ω, *r*_o_ and *α*. For ARSLIP, one needs to add another in *γ*_*a*_.

#### 2.1.3 ARSLIP expands the parameter space over which biological gaits are possible

To determine the gaitspace, MatLab simulations were performed using the non-dimensional equations (2.9). Every parameter set that satisfied the constraints set out in height ratio (2.1), velocity ratio (2.2) and VGRF ratio (2.3) and VGRF profile (2.4) gave rise to, according to our definition, a realistic gait; for these gaits, we also obtained the Froude number,

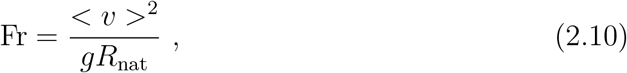

that characterizes the walking speed, < *v* > being the average walking speed over the step.

Figure 2A shows the region where constraints (2.1–2.4) is satisfied. Figure 2A was constructed by combining a series of two dimensional slices each corresponding to different values of *α*. Figure 2A uses *r*_o_ = 0.95, a value close to that observed in humans (see section 2.2.5). The effect of changing *r*_o_ is shown in Supplementary Figures 1 and 2; the results are similar for different *r*_o_.

**Figure 2:**
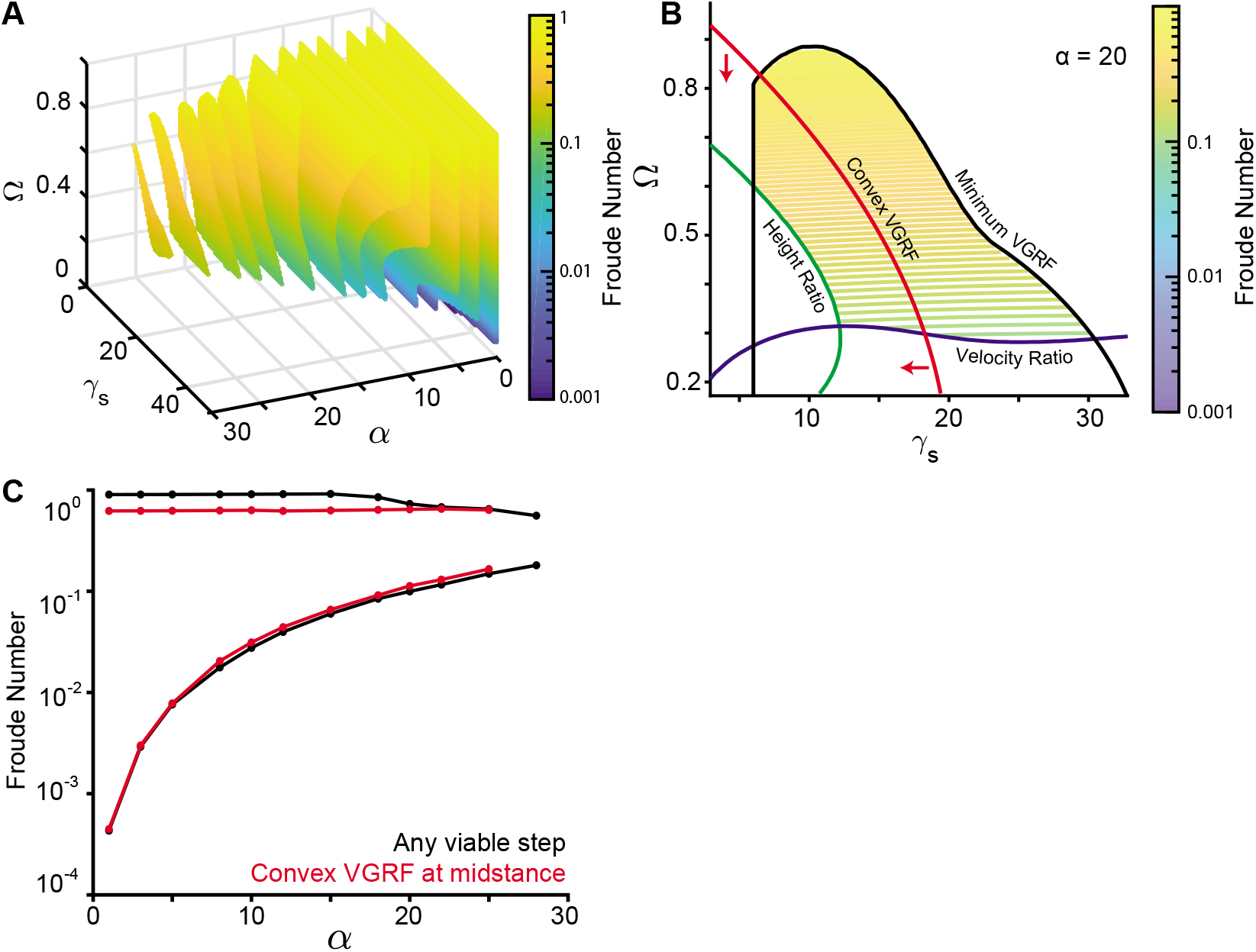
Single SLIP can produce M-shaped vertical GRFs. **A.** Sequence of 2D color-contour plots depicting the froude number for the color contour plot for the allowed gait parameters *γ*_*s*_, *α*, and Ω. Each slice corresponds to one value of *α*, while the vertical midstance height *r*_*o*_ is fixed at 0.95. **B.** 2D contour plot of the *α* = 20 degree slice for *r*_*o*_ = 0.95 as in A. Black represents the VGRF criterion boundary, blue the velocity criterion boundary, and green the height criterion boundary. The red line separates the concave and the convex VGRF regions, with the convex region where VGRF is minimum denoted by the red arrows. **C.** This depicts the minimum and maximum Froude numbers for the gaits that meet the Vr, Hr, and Gr constraints in solid black, while the red line represents additionally meeting the convex VGRF criterion. For each *α, γ*_*s*_ and Ω were varied to obtain the slowest and fastest steps. *r*_*o*_ was fixed to 0.95 like in A.

Fig. 2A ishows that as *α* increases the SLIP gaitspace shrinks, decreasing the range of Fr’s allowed. The reason for this decrease is shown in Fig. 2B where the two dimensional slice for *α* = 20°, an angle that is typical for humans, is plotted. Slow speeds below Fr =0.1 are not allowed. In SLIP there has to be an approximate balance between the radially outward centrifugal and tension forces on one side, and the radially inward component of gravity on the other. For slow speeds since Ω that controls centrifugal force is small, *γ*_*s*_ that is responsible for the spring force, must be large. However, heuristically one expects the oscillation time, 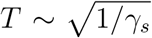, and thus if *γ*_*s*_ is too large, the leg relaxes to its natural length too quickly for the leg to sweep through the given stance angle. In other words, unless *α* is appropriately small, SLIP cannot account for slow speeds. This is indeed consistent with previous findings in [5,28]. Fig. 2C shows the range of allowed speeds as a function of *α* where all other parameters were varied to obtain the maximum and minimum allowed speeds.

The fact that a single springy leg can produce the M-shaped GRF is somewhat surprising from the perspective of previous work done on DSLIP and its variant where the peaks of the M-shape invariably occur during the double support phase [19, 38], instead of during the single stance phase. This ability of SLIP is critical to its appropriateness as a model for walking because empirical GRF peaks are invariably during the single-stance phase [12, 26, 10, 28].

The gaitspace for ARSLIP is shown in Figure 3. Fig. 3A and B show the same plot as Fig. 2A and B, except that now the additional parameter, *γ*_*a*_ is set at 0.5; equivalent plots for other values of *γ*_*a*_ are in Supplement D. Fr can reach lower values; for instance, the lowest Fr in Fig. 3B is about 0.02 whereas in Fig. 2B it is about 0.28. As one increases *γ*_*a*_ it becomes possible to achieve progressively slower speeds (Fig. 3C). As noted in [5], one way to see this effect is to consider the small angle limit of ARSLIP evolution, |*ψ*| ≪ 1 and ignore the slight variations in *r*, so that the evolution equation for *ψ* becomes

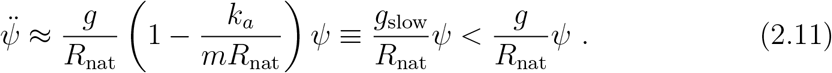

We note that 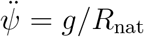, is the equation for an inverted pendulum rotating under the influence of gravity. (2.11) shows that angular springs can counter gravity by effectively making it weaker.

**Figure 3:**
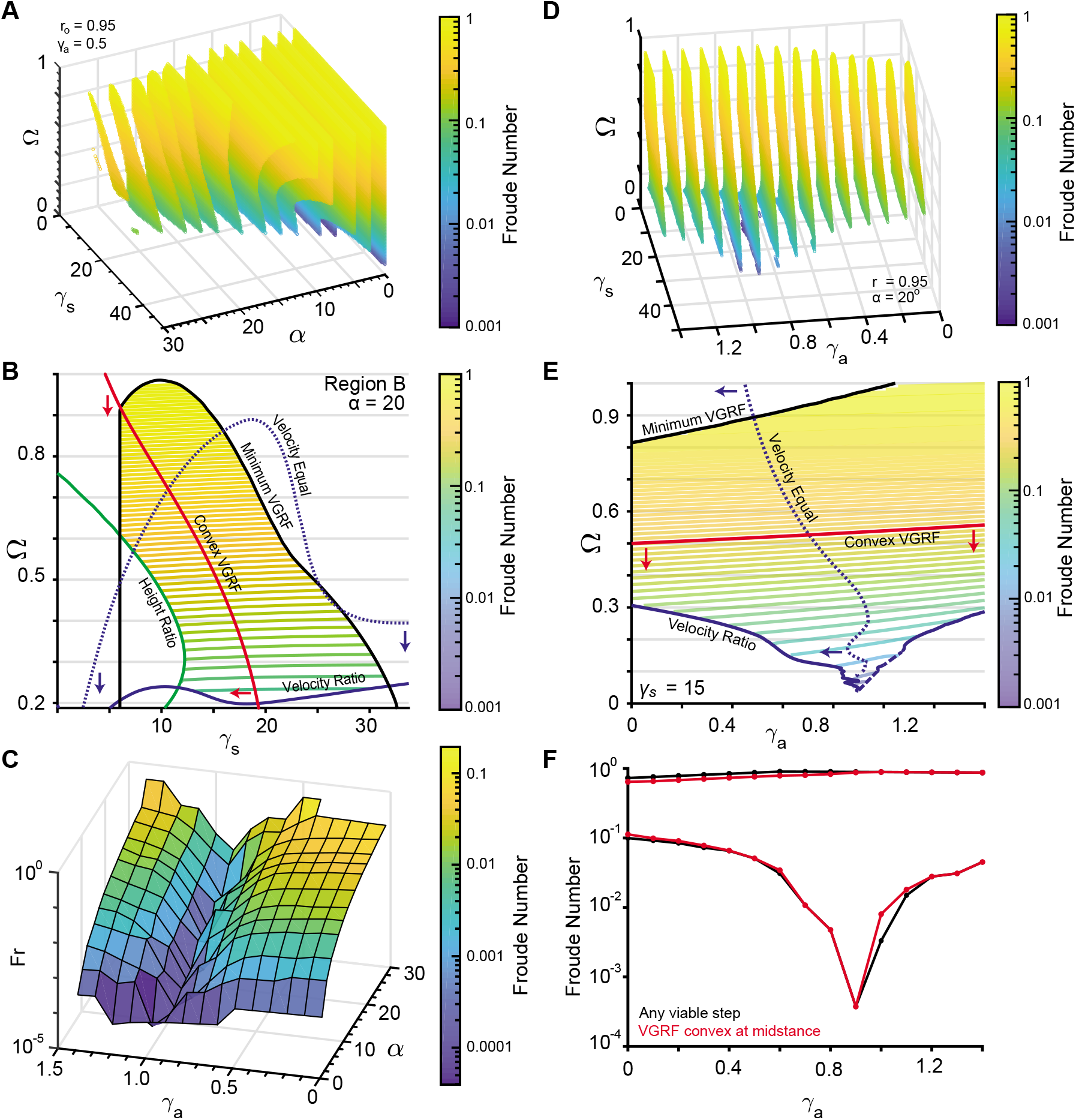
The angular spring in ARSLIP allows for modelling much slower locomotion. **A.** A sequence of 2D color-contour plots depicting the Froude number for the color contour plot for the allowed gait parameters *γ*_*s*_, *α*, and Ω with the vertical midstance height *r*_*o*_ is fixed at 0.95 and *γ*_*a*_ set to 0.5. **B.** 2D contour plot of the *α* = 20 degrees slice for *r*_*o*_ = 0.95 and *γ*_*a*_ *=* 0.5 as in panel A. Similar to 2B, red is the boundary of the minimum VGRF criterion, blue the velocity criterion, and green the height criterion. The dotted line represents the region where the velocity at end of stance equals the midstance velocity. Red arrows indiciate the region where the vertical ground reaction force is minimum, while blue arrows point to the region where the velocity at midstance is less than the velocity at the end of single support. Contour lines represent the Froude number of the allowed gaits. **C.** This depicts the minimum Froude numbers for the gaits that meet the Vr, Hr, and Gr constraints, plotted against *α* and *γ*_*a*_ with a fixed *r*_*o*_ of 0.95 and minimizing over the other parameters. As *γ*_*a*_ increases, the slowest steps that can be modeled decrease. **D.** 2D color-contour plots with color representing the Froude number with Ω, *γ*_*a*_, *γ*_*s*_ varying, and *r*_*o*_ set to 0.95 and *α* to 20 degrees. Each slice corresponds to different values of the angular spring. **E.** 2D slice from D, with *γ*_*s*_ = 15. Same coloring as panel B. The dashed line along the velocity ratio criterion indicates that the boundary is linearly interpolated since the simulation could not resolve the boundary continuously. **F.** The range of Froude numbers possible when *α* = 20 and *r*_*o*_ = 0.95 for different values of *γ*_*a*_ with *γ*_*s*_ and Ω free to vary. The black curve represents any steps that meet Vr, Hr, and Gr, while the red curve is for the steps that additionally meet the convex VGRF criterion.

The effect of the angular spring on gaitspace is shown in Fig. 3D. At a given value of *α*, the gaitspace expands till it reaches a critical value. A similar trend is observed at fixed value of *γ*_*s*_ (Fig. 3E). This critical value does depend on *α* as can be seen in Fig. 3C; for smaller *α* the critical *γ*_*a*_ is also small. The range of Fr accessible in ARSLIP as a function of *α* is shown in Fig. 3F. This range is much larger than the range in SLIP Fig. 2C. ARSLIP, surprisingly, also improves the upperbound because the tangential forces are able to provide greater control so that the speed, and hence the centrifugal force, doesn’t increase as much as in SLIP when the leg rotates away from the midstance. This implies that the VGRF can be slightly larger in ARSLIP as compared to SLIP and the constraint (2.3) is easier to satisfy.

The ability of ARSLIP to produce smooth COM kinematics can equivalently be described in terms of force. In SLIP (or IP for that matter), the leg spring that counteracts gravity also provides a destabilizing horizontal component that speeds up the animal as it moves forward from the midstance. In ARSLIP, the angular spring provides an opposing horizontal component that allows an individual to control its increase in speed. Thus our analysis predicts that the HGRFs in ARSLIP will be smaller than in SLIP and this may address the deficiency that HGRF amplitudes observed in SLIP fits of human GRFs [30] are larger than those observed empirically.

Under some parameter conditions the horizontal component of the angular spring is larger in magnitude than its radial spring counterpart causing the leg to decelerate from its midstance position (“inverted gait”). The gaitspace to the left of the black curve in Figs. 3B and 3E corresponds to the regular gait where the speed increases as the animal goes from the midstance to the beginning/end, while the region on the right correspond to the inverted gait. The inverted gait is observed in some animals such as stick insect and fruit flies [21, 34].

### 2.2 Fitting models to GRFs during walking: SLIP can model the VGRF, but needs ARSLIP to model HGRF

The non-dimensional analysis in the previous section, consistent with findings in [30, 25], show that it is challenging for SLIP to model slow uniform walks. The analysis also shows (see Fig. 3C) that there are two ways to address this issue: Either the subjects can reduce the sweep angle, *α*, as they decrease speed in which case SLIP could continue to be a decent model (Figs. 2A and C). The alternative would be to introduce tangential forces, or *γ*_*a*_, as in ARSLIP that allows for greater speed control. To distinguish between these possibilities and to assess the general effectiveness of SLIP and ARSLIP models in describing realistic walking dynamics, we fit the single stance phase of human walking to these models.

#### 2.2.1 Obtaining Experimental Data

Data was acquired as described previously [23]. Briefly, eight subjects (mass: 74 ± 18 kg, height: 1.69 ± 0.12 m, age: 24 ± 4) walked across four Bertec force plates at a self-selected speed, 20% faster than self-selected speed, 20% slower than self-selected speed, and 50% slower than self-selected speed. The force plates were sampled at 1000 Hz and arranged such that the force readings from the left and right leg were each sampled by two force plates each during experimental trials. The coordinate positions of 47 reflective markers on the subject were recorded using the VICON system sampling at 200 Hz. The COM position of each subject during the trial could be calculated from these markers. Single support phases were extracted from the data, see Fig. 4. Runs in which the single stance phase could not be extracted from the data were removed. Effective leg length was calculated as the distance of the COM to the point underneath the COM at mid-stance. The angle of the leg is the angle the COM position makes with the vertical. For each single stance phase, the COM motion was interpolated using a piecewise cubic hermite interpolant such that for each ground reaction force sample there would be a corresponding COM position.

**Figure 4:**
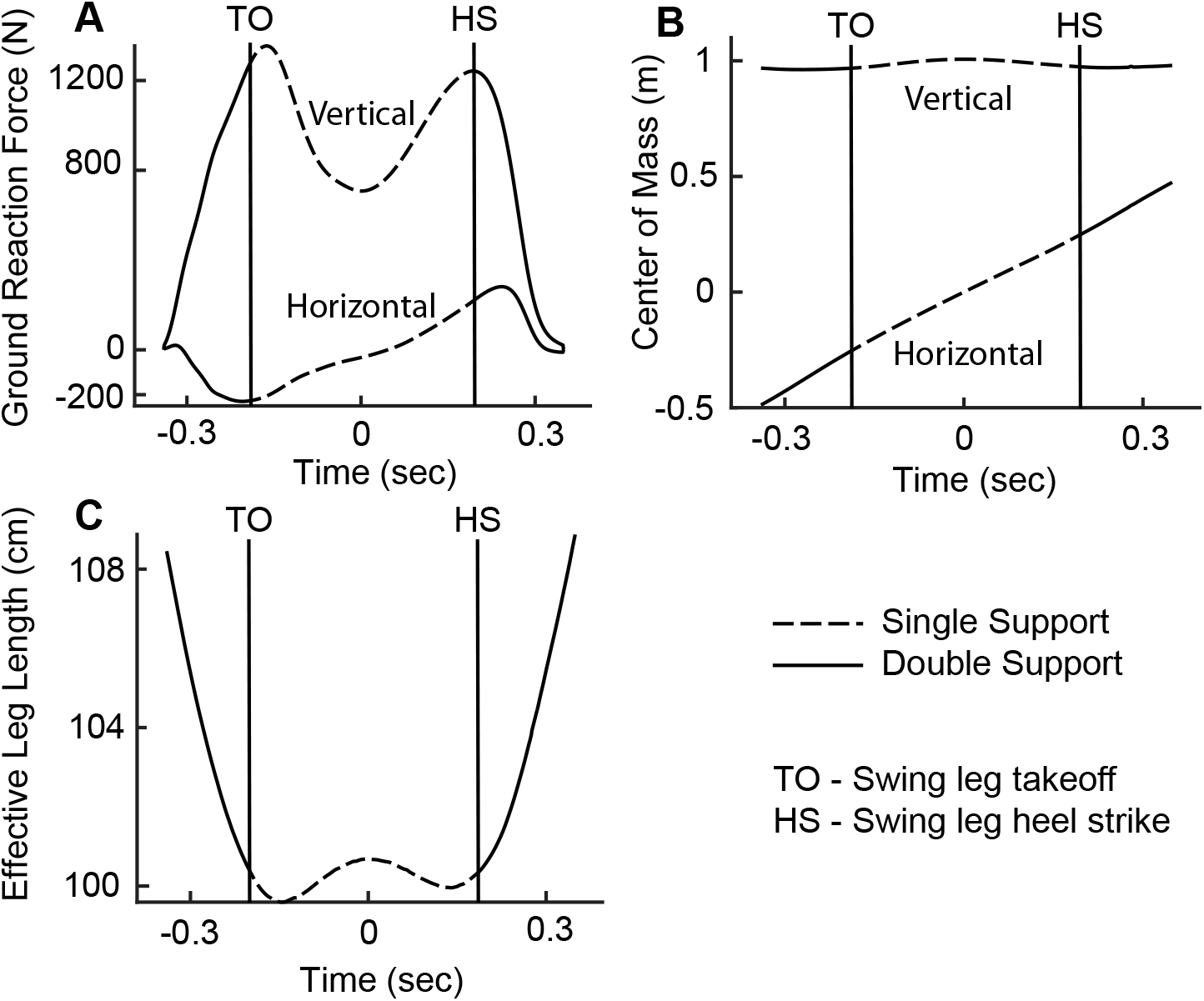
Single support data extracted from the experimental trials in preparation for fitting. **A.** Example of the captured ground reaction forces, with the characteristic M-shaped vertical ground reaction force, and the breaking and then propulsive horizontal ground reaction force. Dashed lines represent the single support phase. **B.** Example of the captures center of mass trajectory. The COM rises towards the midstance and falls away from midstance. **C.** The effective leg length as measured from the center of mass to the projection of the COM at midstance to the ground. The compression as the stance leg is loaded during the beginning of the stance phase is followed by a slight decompression towards midstance. The leg compresses after midstance before unloading in preparation for the swing phase.

#### 2.2.2 Theoretical fits to Experimental Data

A Matlab program using the global search algorithm from the Global Optimization Toolbox was written to fit the parameters of the governing differential equations of SLIP and ARSLIP (2.5 & 2.6) to each single stance phase, minimizing the root mean square error of the predicted GRFs and the sampled GRFs. The leg spring stiffness, *k*_*s*_, was constrained to lie between 0 and 50,000 N/m while the angular spring stiffness, *k*_*a*_, could vary from 0 to 5000 Nm/rad. The natural uncompressed leg length, *R*_nat_, was constrained to lie within 10% of the midstance leg length. The midstance leg length, *R*_o_ and angular speed, *ω*, were constrained by the COM data to vary no more than 2.5% and 5% respectively from their experimental values. Symmetry around mid-stance (*t* = 0) was assumed in the parameter search such that *ψ*(0) = 0 and *ṙ*(0) = 0. Finally, the stance duration was fixed to the empirically observed duration. The best fit parameter was found through gradient descent. Local minimas were avoided by sampling 1000 points within the allowed region. Ignoring points that fell within the same basin of attraction help ensure that the search did not get stuck in a local minima. GRFs were chosen for the objective function over the COM data because of the former’s small experimental error, measured in the tenths of a percent. The COM data was used to constrain the possible leg lengths.

As noted in the introduction, our approach differs from others in constraining force, kinematics and stance duration: In some studies such as in the virtual pivot models [31], only GRFs are considered leading to model parameters that are in clear conflict with the animal’s COM motion as no restriction is placed on the height of the COM. Other studies normalize time to stance duration [10], allow stance time to vary [30], or only focus on fitting the COM trajectory (vertical height as a function of the horizontal displacement) [32].

There are two insights that we obtained from fitting the SLIP and ARSLIP models to human walking data. These insights concern the VGRFs and HGRFs respectively and are discussed in turn in the next two sections.

#### 2.2.3 M-shaped VGRF is caused by CECE of the leg, and to a first approximation, explained by single SLIP

First, we noticed that in many steps, the M-shape of the GRF occurs within the single-stance phase (Figure 5A). Across all the steps, the first peak of the VGRF occurs within the single stance phase, and the second peak occurs right at toe-off (Figure 5E). Importantly, the leg length has a W-shaped profile (Figure 5A) during the single stance phase. Therefore, the M-shaped VGRF can be modeled through spring-like forces resulting from compression and extension of a single leg spring. Specifically, the effective leg length undergoes initial compression as the leg is loaded (causing peak VGRFs) and then decompresses (VGRF minima) as it approaches the mid-stance. After the mid-stance the effective length leg retraces its path leading to an almost symmetric evolution around the mid-stance. As a result of the decompression around midstance, the COM height is at its maxima and the VGRF is at its minima at the mid-stance, a quintessential characteristic of human walking. This VGRF profile can be fit to SLIP; the fitted model also goes through a CECE cycle (5A-C). ARSLIP provided a small, but significant improvement to the VGRF fit. For the VGRF, the ARSLIP RMS error is 30.7 N (4.1% of maximum VGRF magnitude), while for SLIP it is 35.4 N (4.4%). Fits to each step in our dataset is shown in Supplementary Figures 3, 4, 5.

**Figure 5:**
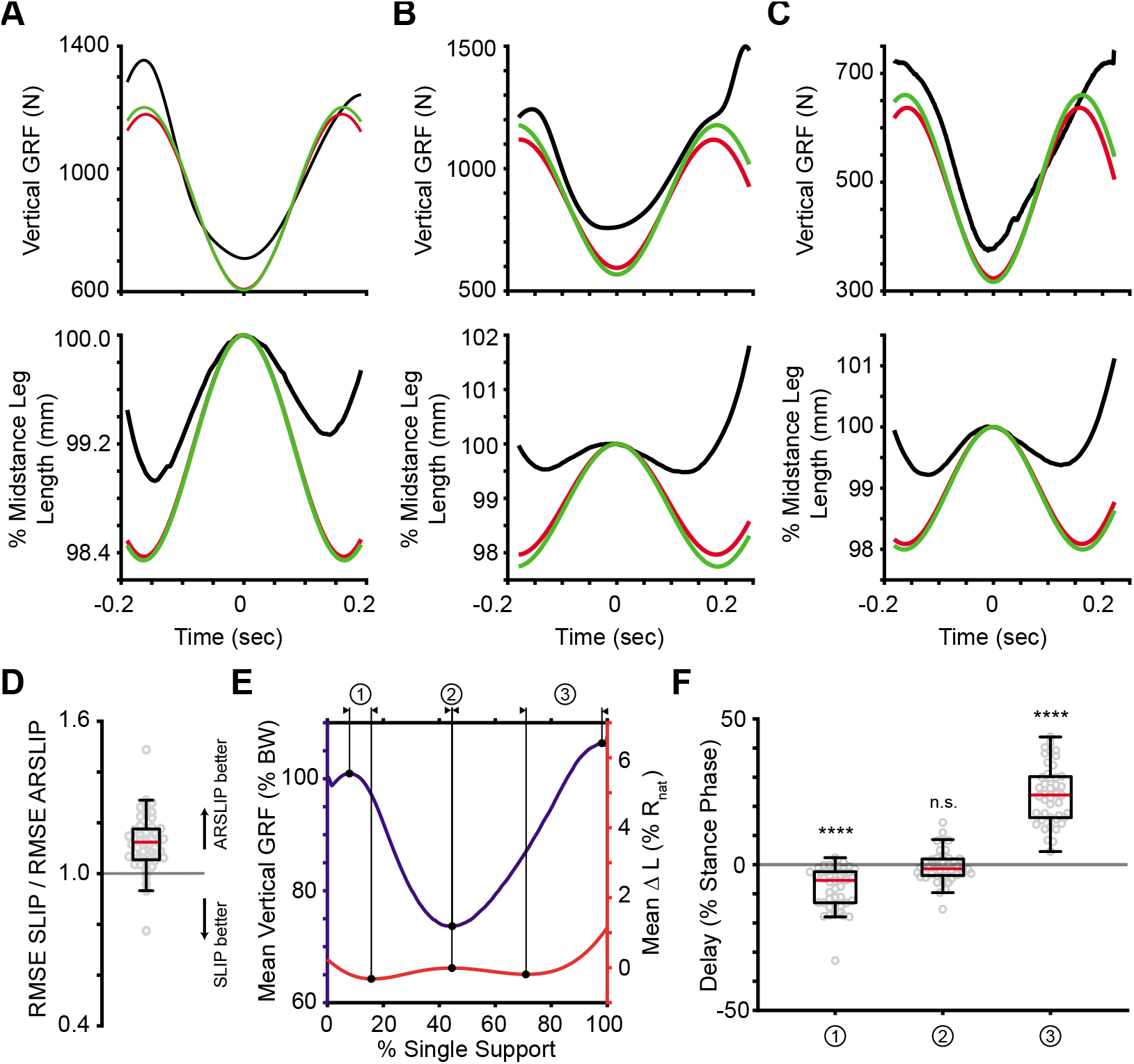
The M-shaped vertical ground reaction force is due to the leg spring of SLIP. **A, B, C** Three example fits from the data showing the relationship between the leg length and the vertical ground reaction force. The change in length of the leg spring drives the M-shaped ground reaction force **D** While both SLIP and ARSLIP result in similar predictions to the VGRF, ARSLIP slightly outperforms SLIP in fitting the vertical GRF. **E** The mean vertical GRF and change in leg length for all trials with each trial normalized to its single support time. Peak vertical GRF are not aligned well with the minimum leg lengths. **F** The delays between the vertical GRF and the change in leg length for each trial as represented by the labels above E. The first VGRF peak comes before the minimum leg length, while the second VGRF peak comes after the minimum leg length. The minimum VGRF occurs during the local maximum in leg length.

The fits also reveal a role for active components. The changes in leg length does not match the changes in VGRF. This mismatch can be seen in individual examples 5B-C, as well as, in the average. The plot of average VGRF and leg length changes demonstrate that the change in leg length lags the changes in VGRF early in the stance cycle, and leads during the late phase of the stance 5E-F.

The large compression of the leg as a person transitions from double stance to single stance (see Fig. 2 in [29] for example) dominates the length changes of a leg during walking. If one considers the entire stance phase, the leg length is near its minima during the entire single stance phase. However, if we consider only the single stance phase, the leg length is at a maxima at mid-stance and compresses on either side of this minima. Both the two minima and maxima occur during the single stance phase and are partly responsible for the M-shaped GRF. We will show later that this CECE behavior of the leg is important for increasing the distance that the body travels during stance, and for decreasing the velocity redirection during the transition.

#### 2.2.4 ARSLIP is a much better fit for HGRF than is the SLIP model

The horizontal GRF stays fairly linear during the single stance phase. However, as predicted from the non-dimensional analysis, with no angular spring, the SLIP leg significantly overestimates the magnitude of the HGRF Fig. 6. Representative examples are shown in Fig. 6A-C. The HGRF amplitude (HGRF magnitude at the beginning subtracted from HGRF magnitude at the end) predicted by the best fit to the SLIP model is particularly large Fig. 6D. The ARSLIP model slightly underestimates the HGRF amplitude, but is a much closer match than SLIP. The median absolute error in the HGRF amplitude for ARSLIP is 38.9 N (20.0%), while it is 113 N (59.4%) for SLIP. The ARSLIP is a much better fit to the data even when the entire time-course is compared. The median root mean square (RMS) error in HGRF for ARSLIP is 10.1 N (9.15% of maximum HGRF magnitude), while it is 38.8 N for SLIP (35.9%). In Fig. 6D and 6D, we have plotted the ratios of the RMS errors of the GRFs from the two model predictions. The addition of the angular spring brings the HGRF more in line with experimental data with a median improvement in the RMS by a large factor of 3.6. Our statistical p-test analysis showed that the probability of SLIP being favored over ARSLIP given the HGRF data fits is less than 0.0001.

**Figure 6:**
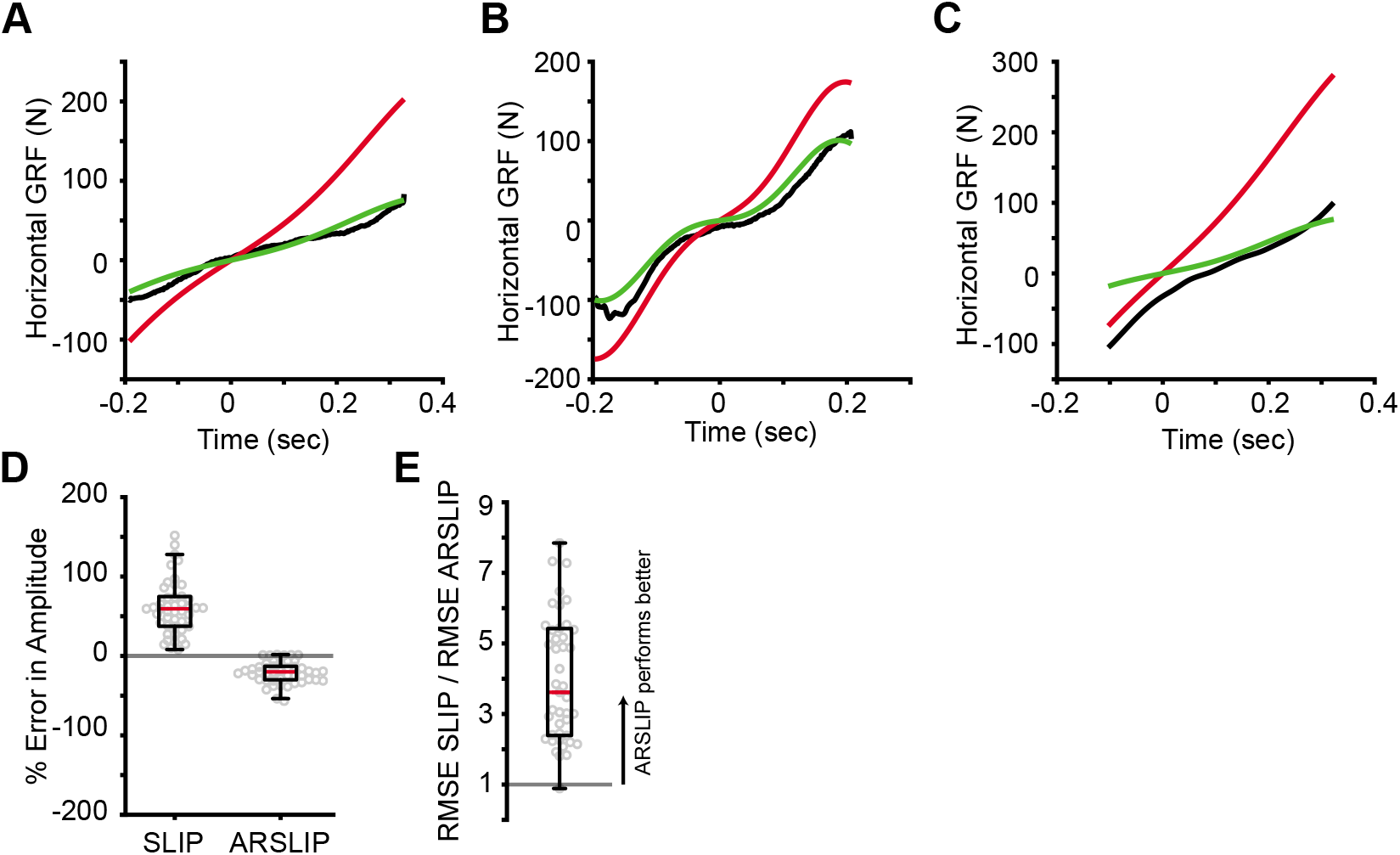
The angular spring improves the fit in the horizontal direction. Three example fits showing **A** SLIP greatly overestimating the horizontal GRF while ARSLIP makes good predictions **B** Both models capable of fitting more complex HGRFs **C** SLIP overestimating and ARSLIP underestimating the HGRF. **D** The error in amplitude of the horizontal and vertical ground reaction forces. ARSLIP tends to underpredict the amplitude while SLIP tends to overpredict. The magnitude of ARSLIP amplitude error is less than the magntiude of SLIP amplitude error (Mann-Whitney U test, *p* << 0.0001). **E** The ratio of RMSE SLIP to RMSE ARSLIP. ARSLIP performs much better than SLIP in the horizontal direction (sign test, p << 0.0001).

#### 2.2.5 Three mechanisms for speed control:step amplitude, radial spring constant and angular spring constant

The non-dimensional analysis suggested that the only way for SLIP to model slow steps is to decrease *α*. Indeed, we observed that while the stance duration increases only slightly with a decrease in the speed, the angle of sweep decreases by nearly a factor of two from the slowest to the fastest step, see Figs. 7A. The slowest steps, Fr ~ 0.05, also had the smallest *α* ~ 11°. At *α* ~ 11°, the minimum Fr that we can estimate from SLIP gaitspace is about 0.02, which shows the consistency between our gaitspace and human data analysis.

**Figure 7:**
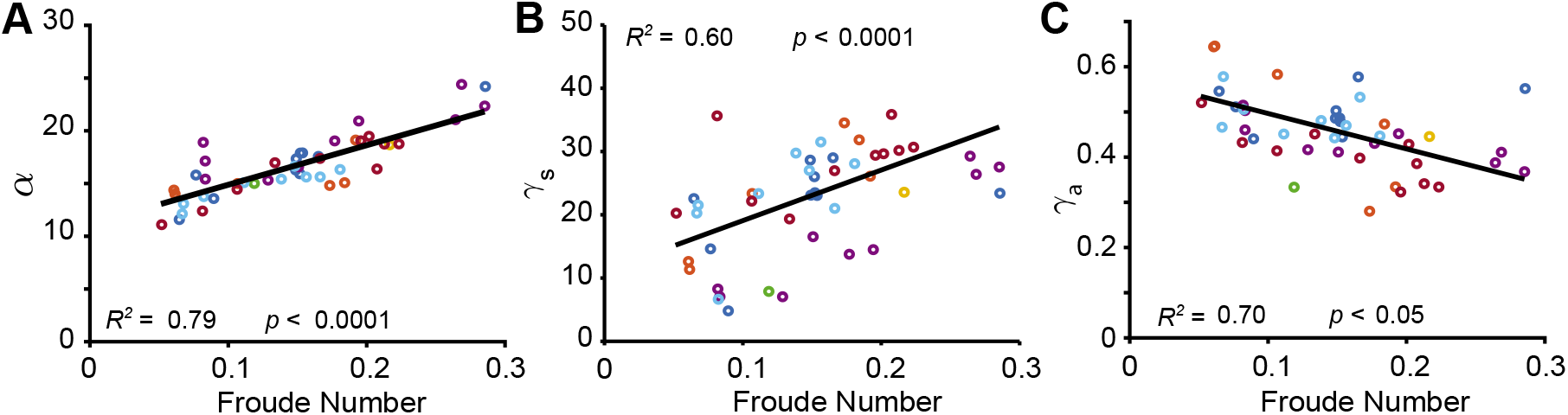
Froude dependent relationships of the dimensionless template parameters and the gaitspace constraints. For each linear plot, the *R*^*2*^ value is for the entire mixed effect model, and the *p* value corresponds to the significance of the fixed effect slope **A.** Maximum angle of sweep **B.** Leg spring **C.** Angular spring

Heuristically speaking, one expects the period of oscillation, *T*, to change as 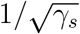, especially for faster motion when the effects of the angular spring is small. And, since < *v* >~ 1/*T*, this implies that *F*_*r*_ ~ *γ*_*s*_. Fig. 7B indeed shows evidence of such an overall linear increase (95% confidence interval for fixed effect slope: [53.77, 106.0]). We note that previous studies [28] also recorded increased *γ*_*s*_ with speed. For discussions on individual variations, please see Supplement F.

Based on our gaitspace analysis we expect *γ*_*a*_ to be increasingly important as the speed decreases. We observe the expected trend in human data. A mixed linear model showed that the angular spring stiffness increases with a decrease in speed. This is consistent with the notion that the angular spring is needed to mitigate unduly large destabilizing effects of gravity on the COM during slower locomotion (see Figure 7C).

### 2.3 Walking in a CECE SLIP cycle contributes to energetically efficient gait

Why do most mammals appear to walk with a M-shaped VGRF? The IP model has a concave VGRF. The radial spring is very stiff, and one would expect its kinetics to approximate that of the IP model. Instead, it generates kinetics that are dramatically different from the IP model. Does being slightly compliant has any biological advantage? In other words, what are the implications of the CECE sequence instead of the CE sequence?

In order to produce a CECE pattern, at the mid-stance the leg length has to be a little longer than that predicted by the fixed point of the system (8A). At low Ω observed during walking, SLIP functions as a stable oscillator along the radial direction (see Supplementary B.2). As a result, the leg oscillates around its fixed point. The mid-stance expanded state implies that the stance phase is lengthened because it is unstable to end the stance phase in a compression. Empirically observed single stance phase lasts more than half a time period (8B), the latter being what one would expect from a CE cycle. A longer step-time increases the step length *α* thereby decreasing the cost of transport by traveling a longer distance using the conservative spring system.

**Figure 8.**
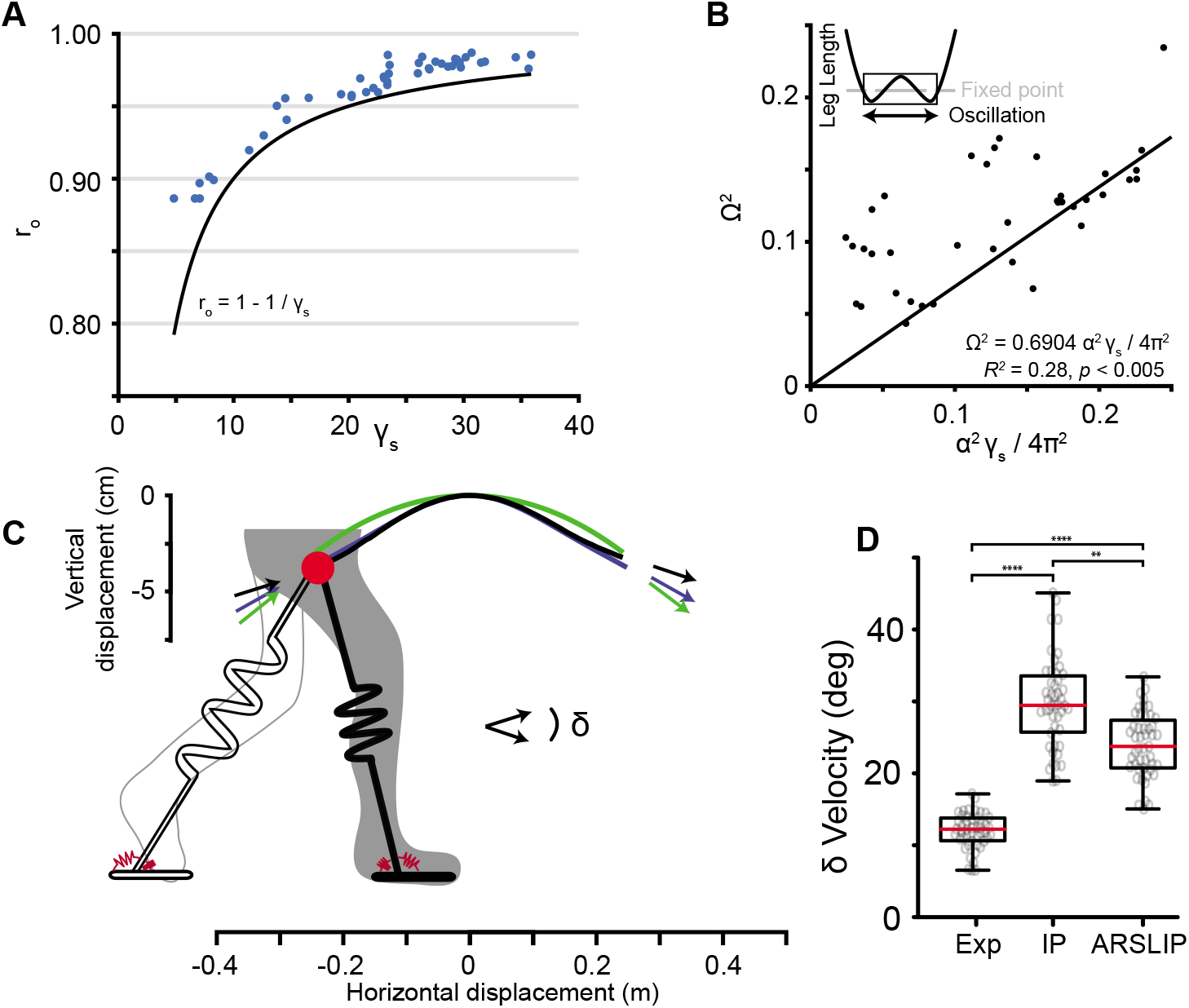
A springy leg reduces the angle the center of mass must be redirected during the double stance phase. **A** Inverse relationship between the leg stiffness and the nondimensional leg length, with all experimental midstance heights (blue dots) above the predicted fixed point of the spring system (black line). **B** About 70% of the natural oscillation of a spring occurs away from midstance during the single support phase. In other words, the single support phase is slightly longer than the natural oscillation period of the leg spring **C** The trajectory of experimental COM (black), ARSLIP fit to COM data (blue) and inverted pendulum (green). The arrows represent the COM velocity vector at the beginning and end of stance. **D** The distribution of the hanges in the velocity angle of the experimental, IP, and ARSLIP cases. There are significant differences between the three distributions (Kruskal-Wallis test, p << 0.0001). The experimental change in the velocity angle is significantly less than what IP predicts (Dunn-Sidak correction, p < 0.0001). The spring simulation predicts a smaller change in the velocity direction than IP (Dunn-Sidak correction, p < 0.005), but greater than what is seen experimentally (Dunn-Sidak correction, p < 0.01)

The CECE walking mode is also more energy efficient in terms of the energy lost during transition. An influential idea in walking is that the single stance phase can be approximated by a passive model such as the inverted pendulum model, and the transition that occurs in the double stance phase is responsible for redirecting the velocity vector [16, 1]. At the end of the stance, COM velocity is directed downwards and has to be redirected upwards at the beginning of the next step. We measured the empirical velocity redirection by assessing the change in velocity vector between the beginning of single stance and the end of single stance. As expected, this change in the velocity vector is small because part of the velocity redirection occurs in the single stance phase. We also computed the velocity redirection necessary if the single stance was modeled by ARSLIP undergoing a CECE cycle or by the IP model. For this purpose we fit ARSLIP and IP to the individual steps but this time minimizing the COM error (8C). Indeed, the velocity redirection necessary was smaller in ARSLIP as compared to IP (8D). The results however may explain why we may prefer a compliant gait as opposed to a the stiff IP gait.

## 3 Discussions & Conclusions

### 3.1 SLIP and ARSLIP as a model for walking: insights and limitations

During most steps (43/47), the leg length features a W-shape during the single support phase, implying that the leg undergoes a CECE cycle. A stiff spring (average stiffness 20kN/m) undergoing a compression of few centimeters can produce the experimentally observed VGRF. The small length changes underlying the W-shape is dwarfed by the large changes in leg length at the transition between single and double stance phases. If the stiff spring of the single-stance phase were to undergo these dramatic length changes, the resulting VGRF changes would be very large. Thus, the spring constant at the transition must be less stiff, and it is unlikely that the full stance phase in humans can be described using a single spring constant. The need for two springs also explains why VGRF fluctuations observed in DSLIP models are usually much larger than empirically observed VGRFs [25].

DSLIP also produces short stance durations. The W-shaped leg length is one mechanism by which stance duration is lengthened. Because the spring undergoes a CECE cycle, the stance duration is lengthened. The CECE cycle also helps in keeping the COM trajectory relatively flat. Another mechanism that helps lengthen the stance duration is the angular spring of the ARSLIP model. The angular spring increases the stance duration by 5-10%.

Where ARSLIP is absolutely necessary is for producing HGRF forces whose magnitude match empirical data. However, because some of the transition from the single support phase to the double support phase occurs during the single support phase, passive models are unlikely to capture the entire dynamics of even the single-support phase of walking. This limitation is apparent in the small mismatch between the timing of the peaks in the GRF and the peaks in the leg length. In future work, adding a simplified transition mechanism to the passive single stance that is described by a stiff spring going through a CECE cycle would further improve the description here, and would likely serve as a complete model for human walking.

### 3.2 SLIP works in a mode which supports energy efficient walking without large changes in COM height

Work started in the late 1950’s in understanding human walking emphasized the importance of walking without large vertical changes in the height of the COM as a mechanism for achieving energy efficiency in walking. Recent work has shown that minimizing vertical movement of the COM does not necessarily result in minimizing work [20, 37]. Inverted pendulum (IP) has a diametrically opposite prediction for energy efficient COM kinematics wherein the COM undergoes large changes in height. However, it is clear that the changes in the vertical height of the COM during walking is much smaller than that predicted by the IP model [29]. The general idea that much of the step - except for the short transition period between one step to the next - can be modeled by a passive model is attractive. We show that employing SLIP as the passive model instead of IP can help some of the inconsistencies above.

An important point to note is that in the context of walking, SLIP functions as a stable system (See Eqn. B.24) that oscillates about the fixed point determined by the system consisting of the leg spring. Importantly, at mid-stance the leg is at its most expanded and the COM is slightly above this fixed point. From this expanded state, the leg contracts before expanding to its original length. There are three consequences of the expanded leg position at mid-stance on the movement of the animal’s COM. First, it increases the duration of the passive phase allowing the animal to travel a longer distance during the passive phase. Second, using a springy mechanism allows much greater control over the speed with which the animal moves through the stance phase. As the animal walks faster, the spring becomes stiffer and the stance duration shorter. Third, the changes in the height of the COM can be smaller than that predicted by the IP model. The change in height of the COM if the leg length is fixed is given *l* sin(*ψ*); as *ψ* increases. The net change in the height of the COM is less because at large angles when a fixed length leg would lose its height, the leg length is increasing, and partially compensates for the decrease in height due to geometry.

We also show that employing a spring as the passive element will also decrease the amount of velocity redirection that needs to be performed at each step. In sum, the flat trajectory proposed by Inman and others, and the passive models proposed later are not as much at odds as it might appear at first glance.

### 3.3 ARSLIP as a general model for locomotion

This study shows that in the context of human walking, the leg spring is stiff. In comparison, the angular spring is relatively weak. Therefore, the angular spring has a relatively small effect on the kinematics in the context of human walking. The most noticeable contribution of the angular spring is in decreasing the HGRF. The angular spring also makes the stance duration longer, but the stance duration is lengthened by only a few percentage points.

However, the angular spring makes a much larger impact on the locomotion of fruit flies. In the case of fruit flies, the angular spring is much stronger in comparative terms, it is strong enough that the acceleration due to the angular spring dominates the deceleration due to the leg spring [13].

More generally, based on whether there is a mid-stance maxima in speed or height, there are 4 kinematic patterns (Figure 9A). All of these kinematic patterns can be produced by the ARSLIP model. These kinematic patterns represent different regimes of the ARSLIP system. It is possible that ARSLIP represents a general model of locomotion.

**Figure 9.**
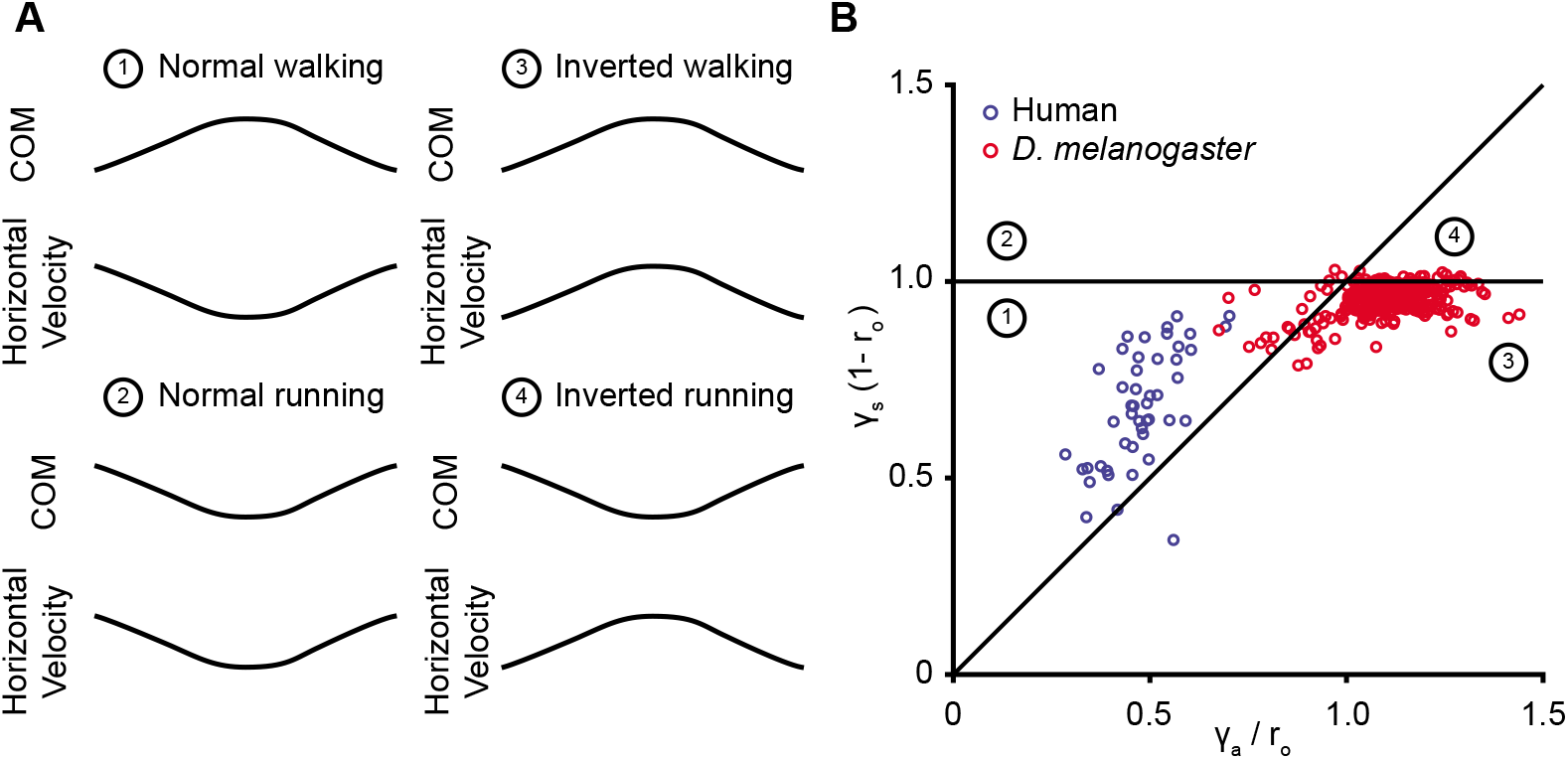
ARSLIP can produce all four kinematic patterns based on COM and horizontal velocity profile. **A** There are four possible locomotion types when looking at COM trajectory and the horizontal velocity profile. The walking gait types have a maximum in height at midstance, while the running a minimum. The inverted locomotion types have a maximum in horizontal velocity at midstance, unlike the minimum that is normally seen. **B** ARSLIP can produce all of these patterns based on the parameters. ARSLIP can capture in one framework the human and fruit fly walking patterns. The numbers correspond to the walking types that were labeled in A.

