## Supplementary material for "Spring-loaded inverted pendulum goes through two contraction-extension cycles during the single stance phase of walking"

### Appendices

#### A Model Formulation and Actualizations

The potential energy of the system, $V$, at any position is the sum of the gravitational and spring potential energies:

$$V = mgR\cos\psi + \frac{1}{2}k_s(R_{\text{nat}} - R)^2 + \frac{1}{2}k_a\psi^2 \tag{A.1}$$

The kinetic energy of the system, $K$, at any position is the sum of the rotational energy of the pendulum and the translation of the COM along the leg as the leg spring compresses:

$$K = \frac{1}{2}m(R^2\dot{\psi}^2 + \dot{R}^2) \tag{A.2}$$

The Lagrangian, $\mathcal{L}$, is defined as $K - V$:

$$\mathcal{L} = \frac{1}{2}m(R^2\dot{\psi}^2 + \dot{R}^2) - mgR\cos\psi - \frac{1}{2}k_s(R_{\text{nat}} - R)^2 - \frac{1}{2}k_a\psi^2 \tag{A.3}$$

$R$ and $\psi$ are the parameters of the Lagrangian. The equations of motion are described by two coupled differential equations:

$$\frac{\partial}{\partial t}\left(\frac{\partial \mathcal{L}}{\partial \dot{\psi}}\right) = \frac{\partial \mathcal{L}}{\partial \psi} \tag{A.4}$$

$$\frac{\partial}{\partial t}\left(\frac{\partial \mathcal{L}}{\partial \dot{R}}\right) = \frac{\partial \mathcal{L}}{\partial R} \tag{A.5}$$

Plugging in $\mathcal{L}$ from Equation A.3 yields

$$2mR\dot{R}\dot{\psi} + mR^2\ddot{\psi} = mgR\sin\psi - k_a\psi \tag{A.6}$$

$$m\ddot{R} = -mg\cos\psi + k_s(R_{\text{nat}} - R) + mR\dot{\psi}^2 \tag{A.7}$$

Rearranging and simplifying the above equation gives the evolution of ARSLIP

$$\ddot{\psi} = \frac{g}{R}\sin\psi - \frac{k_a}{mR^2}\psi - 2\frac{\dot{R}}{R}\dot{\psi} \tag{A.8}$$

$$\ddot{R} = L\dot{\psi}^2 - g\cos\psi + k_s(R_{\text{nat}} - R) \tag{A.9}$$

#### B Derivations for gait criteria

##### B.1 VGRF Concavity

The non-dimensional form of the evolution equations (2.5, 2.6) governing GSLIP are:

$$\ddot{r} = r\dot{\psi}^2 - \cos\psi + \gamma_s(1 - r) \tag{B.10}$$

$$\ddot{\psi} = \frac{1}{r}\sin\psi - \gamma_a\frac{\psi}{r^2} - 2\frac{\dot{r}\dot{\psi}}{r} \tag{B.11}$$

where $r \equiv R/R_{\text{nat}}$, and $\psi$ is the angle of the leg, with $\psi = 0$ at mid-stance.

The equations of motion governing the trajectory of the COM can be expressed in Cartesian coordinates:

$$x = r\sin(\psi) \tag{B.12}$$
$$y = r\cos(\psi) \tag{B.13}$$

where $x$ is the anteroposterior (horizontal) location of the center of mass, $y$ the vertical position of the COM. The acceleration of the COM in the vertical direction is the second derivative of $y$:

$$\dot{y} = D[r\cos(\psi)] \tag{B.14}$$
$$\dot{y} = \dot{r}\cos(\psi) - r\dot{\psi}\sin(\psi) \tag{B.15}$$
$$\ddot{y} = D[\dot{y}] = D[\dot{r}\cos(\psi) - r\dot{\psi}\sin(\psi)] \tag{B.16}$$
$$\ddot{y} = (\ddot{r} - r\dot{\psi}^2)\cos\psi - (2\dot{r}\dot{\psi} + r\ddot{\psi})\sin\psi \tag{B.17}$$

Since the ground reaction force must also account for the weight of the COM, the force acting on the ground becomes:

$$F_y = 1 + (\ddot{r} - r\dot{\psi}^2)\cos\psi - (2\dot{r}\dot{\psi} + r\ddot{\psi})\sin\psi \tag{B.18}$$

where the term 1 is the non-dimensional gravitational force. Substituting equations B.10 and B.11 to the previous equation and simplifying yields:

$$F_y = \gamma_s(1-r)\cos\psi + \frac{\gamma_a\psi}{r}\sin\psi \tag{B.19}$$

$F_y$ is the sum of the forces produced by the leg spring and the angular spring projected on the vertical. Differentiating $F_y$ twice with respect to non-dimensional time, plugging in Equations B.10 and B.11, simplifying, and evaluating at $\psi = 0$ yields

$$\frac{2\Omega^2}{r_\circ}\gamma_a + \gamma_s(1-\Omega^2) - (1-r_\circ)\gamma_s^2 \geq 0 \tag{B.20}$$

for the criterion where the vertical ground reaction force is convex at mid-stance.

#### B.2 Approximate Time Period from Dynamics

Per Equation 2.7, the evolution of the leg length $r$ is:

$$\ddot{r} = r\dot{\psi}^2 - \cos\psi + \gamma_s(1-r) \tag{B.21}$$

Assuming small angles $\psi$ and that the change in angular velocity $\dot{\psi}$ is small, the previous equation reduces to

$$\ddot{r} = r\Omega^2 - 1 + \gamma_s - r\gamma_s = -(\gamma_s - \Omega^2)r - 1 + \gamma_s \tag{B.22}$$

Rearranging the equation leads to the following nonhomogenous equation:

$$\ddot{r} + r(\gamma_s - \Omega^2) = \gamma_s - 1 \tag{B.23}$$

This is analogous to a harmonic oscillator with angular frequency $\sqrt{\gamma_s - \Omega^2}$. For $\gamma_s > \Omega^2$, the solution is stable, and will oscillate around:

$$r^\star = \frac{\gamma_s - 1}{\gamma_s - \Omega^2} \tag{B.24}$$

Because $r^\star$ corresponds to the oscillations of the leg during stance, $r^\star$ must necessarily be less than 1. Because of this, an additional constraint to the solution is required: $\Omega^2 < 1$. Human walking as found in the papers is in the regime of this constraint, as $\gamma_s > 1 > \Omega^2$ for all steps. The oscillation time period of the spring is given by:

$$T_\omega = \frac{2\pi}{\omega} = \frac{2\pi}{\sqrt{\gamma_s - \Omega^2}} \tag{B.25}$$

#### B.3 Requirement of angular spring for inverted gait

There are four possible walking gaits based on the height, $h$, and velocity, $v$, profiles:

1. Regular Walking: $h$ has a maximum at midstance while $v$ has a minimum, like in human walking
2. Inverted Walking: $h$ and $v$ both have a maximum at midstance, like fruit flies and stick insects
3. Regular Running: $h$ and $v$ both have a minimum at midstance, like human or cockroach running
4. Inverted Running: $h$ has a minimum at midstance while $v$ has a maximum, which is unlike any organism that we can recall

In ARSLIP, the boundary between walking and running is given by the hypersurface

$$\ddot{h} = 0, \tag{B.26}$$

in the gaitspace as $\dot{h} = 0$ at midstance due to the symmetric constraint. From Newton's Laws, the force at midstance in nondimensional terms is

$$\ddot{h} = \gamma_s(1 - r) - 1, \tag{B.27}$$

which combined with B.26 implies:

$$\gamma_s = \frac{1}{1-r_\circ} \iff r_\circ = 1 - \frac{1}{\gamma_s} \equiv r_c \tag{B.28}$$

If $r_\circ > r_c$, then $\ddot{h} < 0$, which corresponds to a walking COM profile with a maximum height at midstance. If $r_\circ < r_c$, then $\ddot{h} > 0$, which corresponds to a running COM profile with a minimum height at midstance.

The transition boundary between regular vs inverted velocity profiles is given by the hypersurface where

$$\ddot{v} = 0, \tag{B.29}$$

as $\dot{v} = 0$ for symmetric ARSLIP. To find the transition boundary, we can look at the horizontal GRF

$$\ddot{x} = -\gamma_s(1-r)\sin\psi + \gamma_a \frac{\psi\cos\psi}{r} \tag{B.30}$$

Since $\psi = 0$ at midstance, the right hand side vanishes, implying that $v$ must be a minimum or maximum. Since $\ddot{v} = \dddot{x}$, we can differentiate B.30:

$$\ddot{v} = \dddot{x} = \gamma_s \left[\dot{r}\sin\psi - (1-r)\dot{\psi}\cos\psi\right] + \gamma_a \left[(\cos\psi - \psi\sin\psi)\frac{\dot{\psi}}{r} - \frac{\dot{r}\psi\cos\psi}{r^2}\right] \tag{B.31}$$

Plugging in the midstance values of $\psi = 0$, $\dot{\psi} = \Omega$, and $r = r_\circ$ yields

$$\ddot{v} = \left[\gamma_s(1-r_\circ) - \frac{\gamma_a}{r_\circ}\right]\Omega \tag{B.32}$$

If $\gamma_a = 0$ as is the case for SLIP, the gait is guaranteed to have the regular velocity profile where $v$ has a local minimum at midstance. However, if $\gamma_a > \gamma_s r_\circ(1-r_\circ)$, then one can have the inverted gait.

Combining B.28 and B.32, the following demarcation exists for the different gaits:

$$1 > \gamma_s(1-r_\circ) > \frac{\gamma_a}{r_\circ} \qquad \text{for regular walking}$$

$$\min\left\{1, \frac{\gamma_a}{r_\circ}\right\} > \gamma_s(1-r_\circ) \qquad \text{for inverted walking}$$

$$\max\left\{1, \frac{\gamma_a}{r_\circ}\right\} < \gamma_s(1-r_\circ) \qquad \text{for regular running}$$

$$1 < \gamma_s(1-r_\circ) < \frac{\gamma_a}{r_\circ} \qquad \text{for inverted running}$$

### C Supplementary Methods

To adjust for expected variability due to different strategies of walking between different subjects, a mixed linear model was fit for the angle of sweep, $\alpha$, leg spring, $\gamma_s$, and the angular spring, $\gamma_a$ plotted against the Froude number. Only the subjects for whom multiple trials were extracted were used for this analysis. Random effects could influence the intercept only, slope only, both slope and intercept in a correlated manner, or both slope and intercept in an uncorrelated manner. In Wilkinson notation, these equations can be expressed as:

$$y \sim 1 + \mathrm{Fr} \tag{C.33}$$
$$y \sim 1 + \mathrm{Fr} + (1 \mid \mathrm{Subject}) \tag{C.34}$$
$$y \sim 1 + \mathrm{Fr} + (-1 + \mathrm{Fr} \mid \mathrm{Subject}) \tag{C.35}$$
$$y \sim 1 + \mathrm{Fr} + (1 + \mathrm{Fr} \mid \mathrm{Subject}) \tag{C.36}$$
$$y \sim 1 + \mathrm{Fr} + (1 \mid \mathrm{Subject}) + (-1 + \mathrm{Fr} \mid \mathrm{Subject}) \tag{C.37}$$

The model chosen for the regression was the model whose Aikake Information Criterion (AIC) was the lowest. In Figs. 7 A-C, we have plotted the dimensionless parameters, $\alpha$, $\gamma_s$, and $\gamma_a$ against the Froude number obtained from the fits to the ARSLIP model. The parameter values that we obtain from SLIP fits for the common parameters $\Omega, \gamma_s, r_\circ$ are not that different from ARSLIP values, see Supplement F.

For the angular spring, the model with the smallest AIC was of the form:

$$\gamma_{a_{\mathrm{Subject}}} = \beta_0 + \beta_1 \mathrm{Fr} + \beta_{0_{\mathrm{Subject}}} + \beta_{1_{\mathrm{Subject}}} \mathrm{Fr} \tag{C.38}$$

The model contains a fixed intercept, $\beta_0$, and slope, $\beta_1$, in addition to a random effect for the intercept and slope, $\beta_{0_{\mathrm{Subject}}}$ and $\beta_{1_{\mathrm{Subject}}}$, respectively, which were correlated with one another. The fixed slope is negative, *i.e.* , the angular spring stiffness increases with a decrease in speed (95% confidence interval: $[-1.51, -0.06]$, $p < 0.05$). The mixed effects model could account for 70% of the variability of the angular spring by taking into account the speed and the individual subjects. The This is consistent with the notion that the angular spring is needed to mitigate unduly large destabilizing effects of gravity on the COM during slower locomotion.

For the leg spring, the best fitting model was of the from:

$$\gamma_{s_{\mathrm{Subject}}} = \beta_0 + \beta_1 \mathrm{Fr} + \beta_{0_{\mathrm{Subject}}} \tag{C.39}$$

The $R^2$ value of the mixed effect model was 0.6055. The mean fixed effect slope was 80.48 (95% confidence interval: $[54.358, 106.6]$, $p < 10^{-6}$). As speed increased, the leg spring is predicted to stiffen.

For the angle of sweep, the best fitting model was:

$$\alpha_{\text{Subject}} = \beta_0 + \beta_1 \text{Fr} + \beta_{0_{\text{Subject}}} \tag{C.40}$$

The $R^2$ value of the mixed effect model was 0.7968. The mean fixed effect slope was 37.474 (95% confidence interval: [30.5244.428], $p < 10^{-13}$). As speed increases, the maximum angle swept by the leg is predicted to increase.

### D Effects on Gaitspace from variations in $r_\circ$ and $\alpha$

#### D.1 SLIP

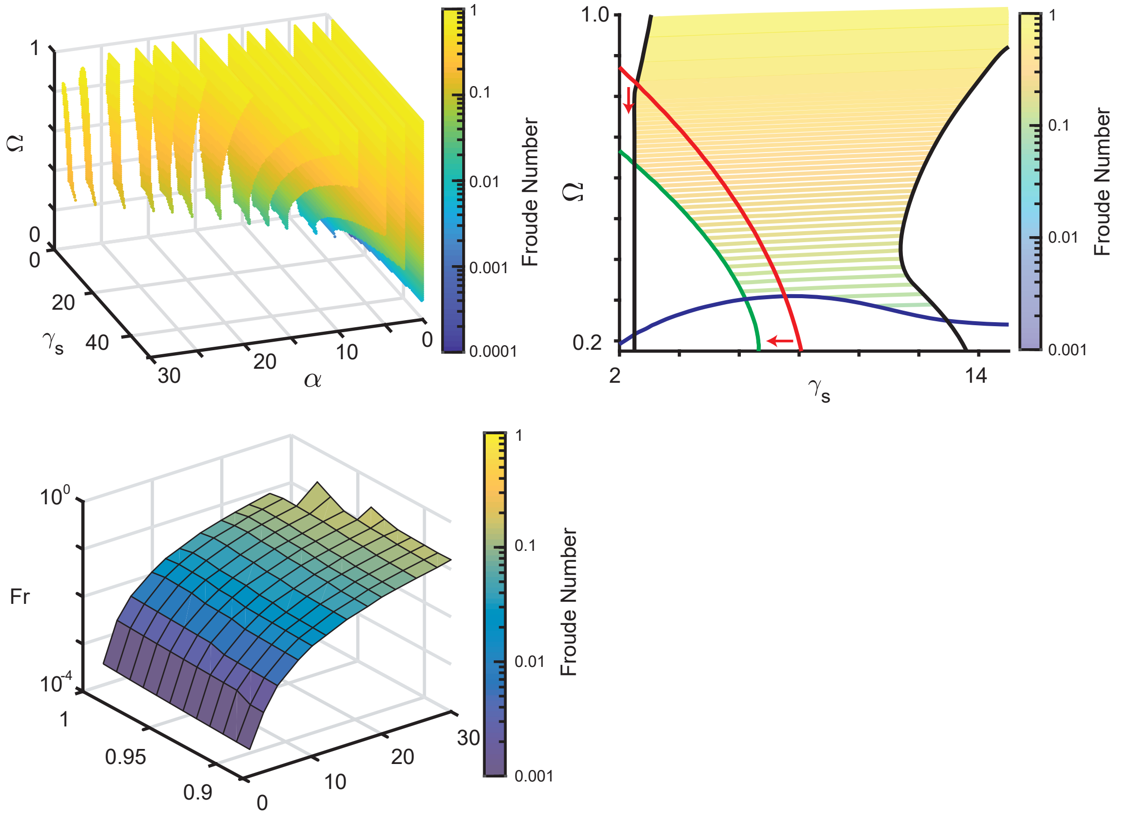


**Supplementary Figure 1: Changing the midstance height does not impact conclusions for analysis of SLIP for modelling of walking. A.** Sequence of 2D color-contour plots depicting the froude number for the color contour plot for the allowed gait parameters $\gamma_s$, $\alpha$, and $\Omega$. Each slice corresponds to one value of $\alpha$, while the vertical midstance height $r_o$ is fixed at 0.88. **B.** 2D contour plot of the $\alpha$ = 20 degree slice for $r_o$ = 0.88 as in A. Black represents the VGRF criterion boundary, blue the velocity criterion boundary, and green the height criterion boundary. The red line separates the concave and the convex VGRF regions, with the convex region where VGRF is minimum denoted by the red arrows. The region for running extends past the simulated region. **C.** 3D surface plot of the minimum allowed Froude values for SLIP when minimizing over $\gamma_s$ and $\Omega$. Note that changing midstance height has little impact on the minimum allowed speed for a given step length.

#### D.2 ARSLIP

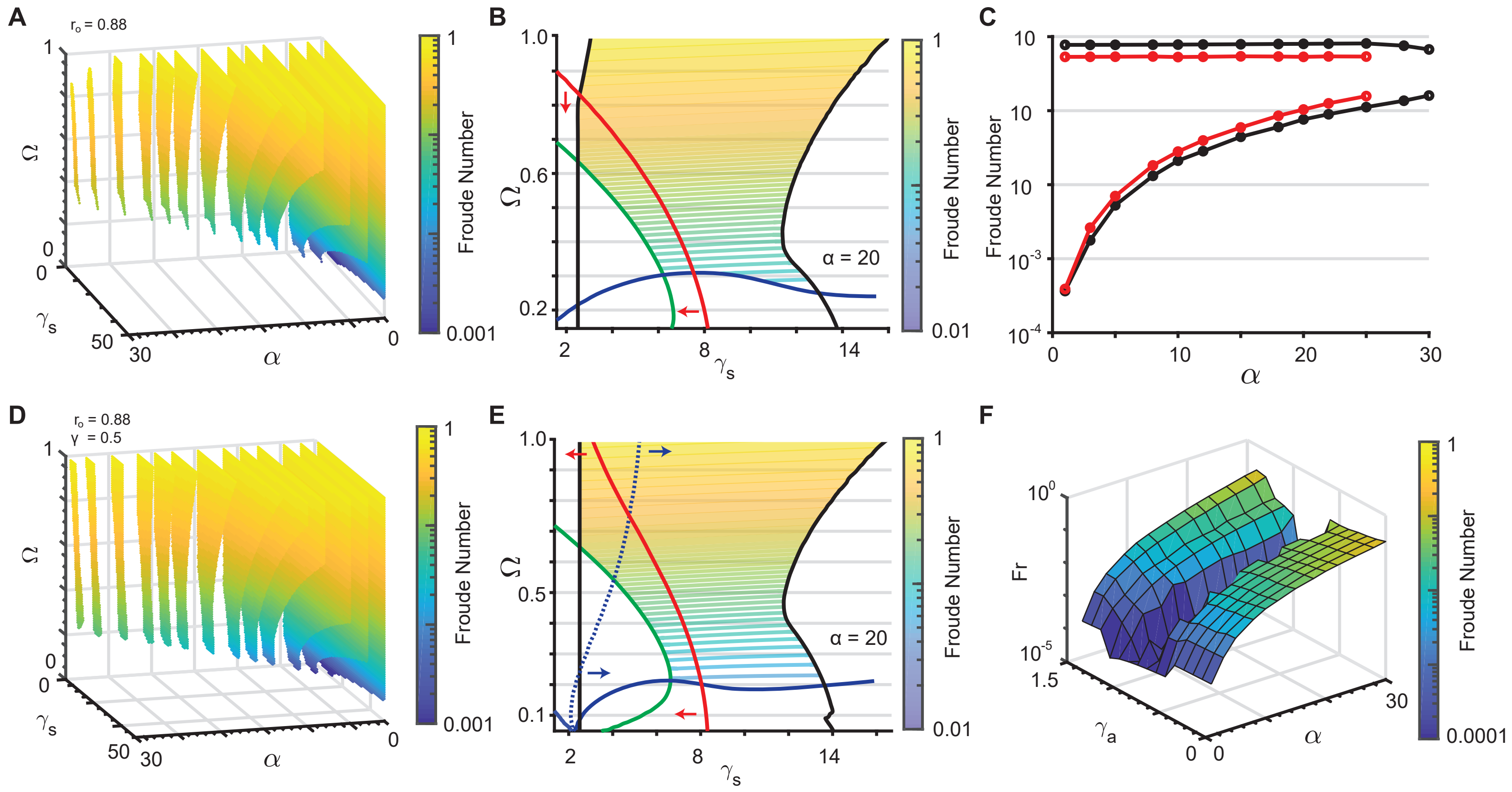


**Supplementary Figure 2: Changing $r_o$ does not influence nondimensional analysis conclusions. A.** Sequence of 2D color-contour plots depicting the froude number for the color contour plot for the allowed gait parameters $\gamma_s$, $\alpha$, and $\Omega$. Each slice corresponds to one value of $\alpha$, while the vertical midstance height $r_o$ is fixed at 0.88. **B.** 2D contour plot of the $\alpha$ = 20 degree slice for $r_o$ = 0.88 as in A. Black represents the minimum VGRF criterion boundary, blue the velocity criterion boundary, and green the height criterion boundary. The red line separates the concave and the convex VGRF regions, with the convex region where VGRF is minimum denoted by the red arrows. **C.** This depicts the minimum and maximum Froude numbers for the gaits that meet the Vr, Hr, and Gr constraints in solid black, while the red line represents additionally meeting the VGRF concavity criterion. For each $\alpha$, $\gamma_s$ and $\Omega$ were varied to obtain the slowest and fastest steps. $r_o$ was fixed to 0.88 like in A and B. **D.** Analogous to A, a sequence of 2D color-contour plots depicting the Froude number for the color contour plot for the allowed gait parameters $\gamma_s$, $\alpha$, and $\Omega$ with the vertical midstance height $r_o$ is fixed at 0.88 and $\gamma_a$ set to 0.5. **E.** 2D contour plot of the $\alpha$ = 20 degrees slice for $r_o$ = 0.88 and $\gamma_a$ = 0.5 as in panel A. Similar to 2B, black is the boundary of the minimum VGRF criterion, blue the velocity criterion, and green the height criterion. The dotted line represents the region where the velocity at end of stance equals the midstance velocity. Red arrows indiciate the region where the vertical ground reaction force is minimum, while blue arrows point to the region where the velocity at midstance is less than the velocity at the end of single support. Contour lines represent the Froude number of the allowed gaits. **F.** This depicts the minimum Froude numbers for the gaits that meet the Vr, Hr, and Gr constraints, plotted against $\alpha$ and $\gamma_a$ with a fixed $r_o$ of 0.88 and minimizing over the other parameters. As $\gamma_a$ increases, the slowest steps that can be modeled decrease.

### E Glossary

| Term | Units | Nondim. | Definition |
|---|---|---|---|
| COM | – | – | Center of mass |
| IP | – | – | Inverted pendulum |
| SLIP | – | – | Spring loaded inverted pendulum |
| DSLIP | – | – | Double SLIP |
| ARSLIP | – | – | Angular and radial spring loaded inverted pendulum |
| GRF(s) | N | – | Ground reaction force(s) |
| HGRF, $F_x$ | N | – | Horizontal ground reaction force |
| VGRF, $F_y$ | N | – | Vertical ground reaction force |
| BW | N | – | Body weight |
| RMSE | – | – | Root mean squared error |
| $R_{\mathrm{nat}}$ | m | 1 | Uncompressed leg length |
| $R_\circ$ | m | $r_\circ = \frac{R_\circ}{R_{\mathrm{nat}}}$ | Midstance leg height |
| $R$ | m | $r = \frac{R}{R_{\mathrm{nat}}}$ | Instantaneous leg length |
| $k_a$ | $\frac{\mathrm{N{\cdot}m}}{\mathrm{rad}}$ | $\gamma_a = \frac{k_a}{mgR_{\mathrm{nat}}}$ | Angular spring stiffness |
| $k_s$ | $\frac{\mathrm{N}}{\mathrm{m}}$ | $\gamma_s = \frac{k_s R_{\mathrm{nat}}}{mg}$ | Leg spring stiffness |
| $\omega$ | $\frac{\mathrm{rad}}{\mathrm{s}}$ | $\Omega = \omega\sqrt{\frac{R_{\mathrm{nat}}}{g}}$ | Angular velocity at midstance |
| $\psi$ | rad | $\psi$ | Angle of leg from vertical |
| $\alpha$ | rad | $\alpha$ | Maximum leg angle from vertical |
| $m$ | kg | 1 | Subject mass |
| $g$ | $\frac{m}{s^2}$ | 1 | Gravitational acceleration |
| $v$ | $\frac{\mathrm{m}}{\mathrm{s}}$ | $\mathrm{Fr} = \frac{v^2}{gR_{\mathrm{nat}}}$ | Froude number |
| $V_r$ | – | – | Velocity ratio (see Fig. 1D) |
| $H_r$ | – | – | Height ratio (see Fig. 1D) |
| $G_r$ | – | – | Ground reaction force ratio (see Fig. 1D) |

### F Model fits to all human stance phase data

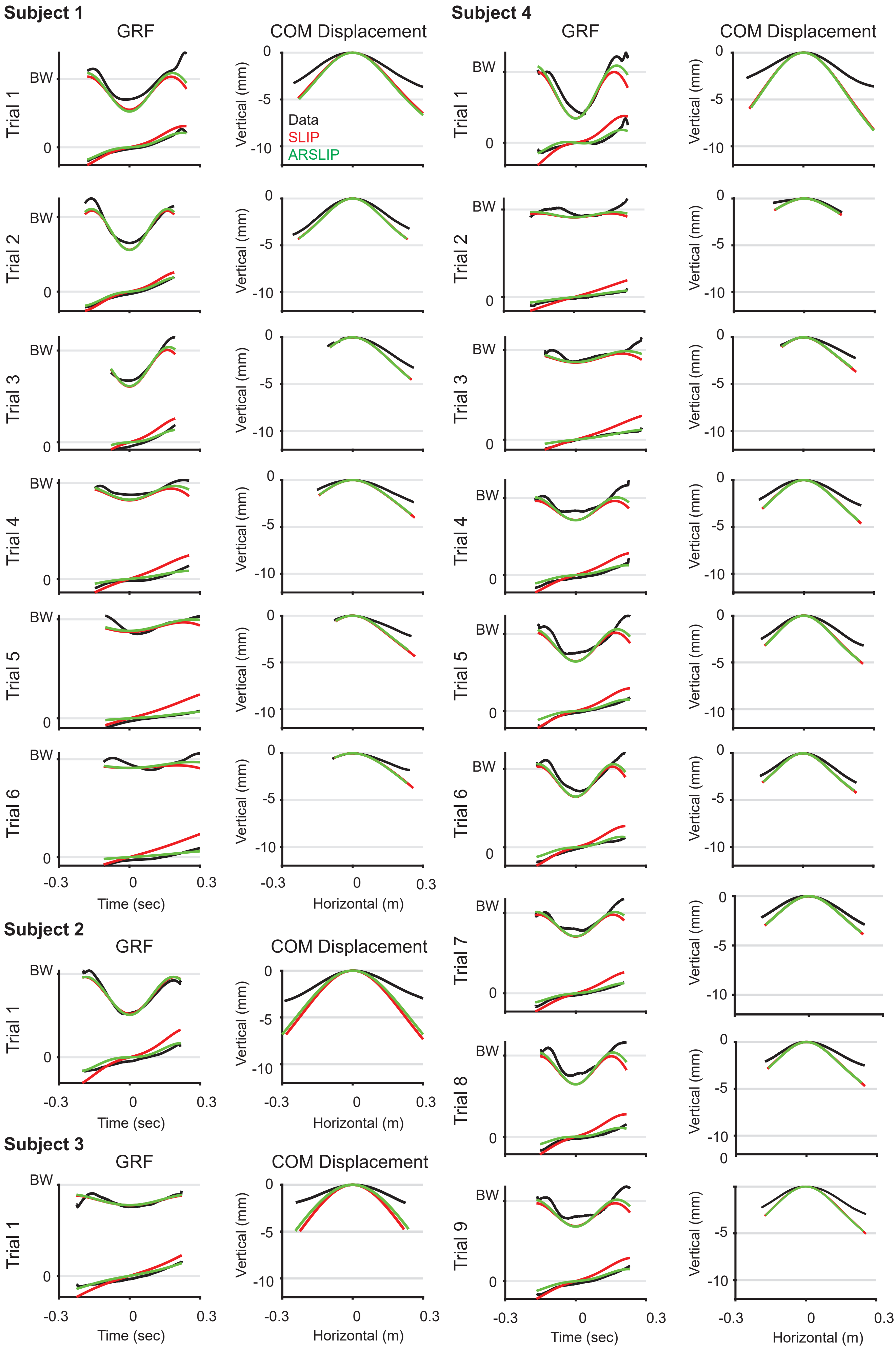


**Supplementary Figure 3: SLIP and ARSLIP fits to individual trials for Subjects 7 and 8.** For each panel, the experimental data is plotted in black, the SLIP fit is in red, and the ARSLIP is in green. For the GRF fits, the GRFs are normalized to subject weight and are plotted versus time. The COM displacement is plotted relative to the midstance position.

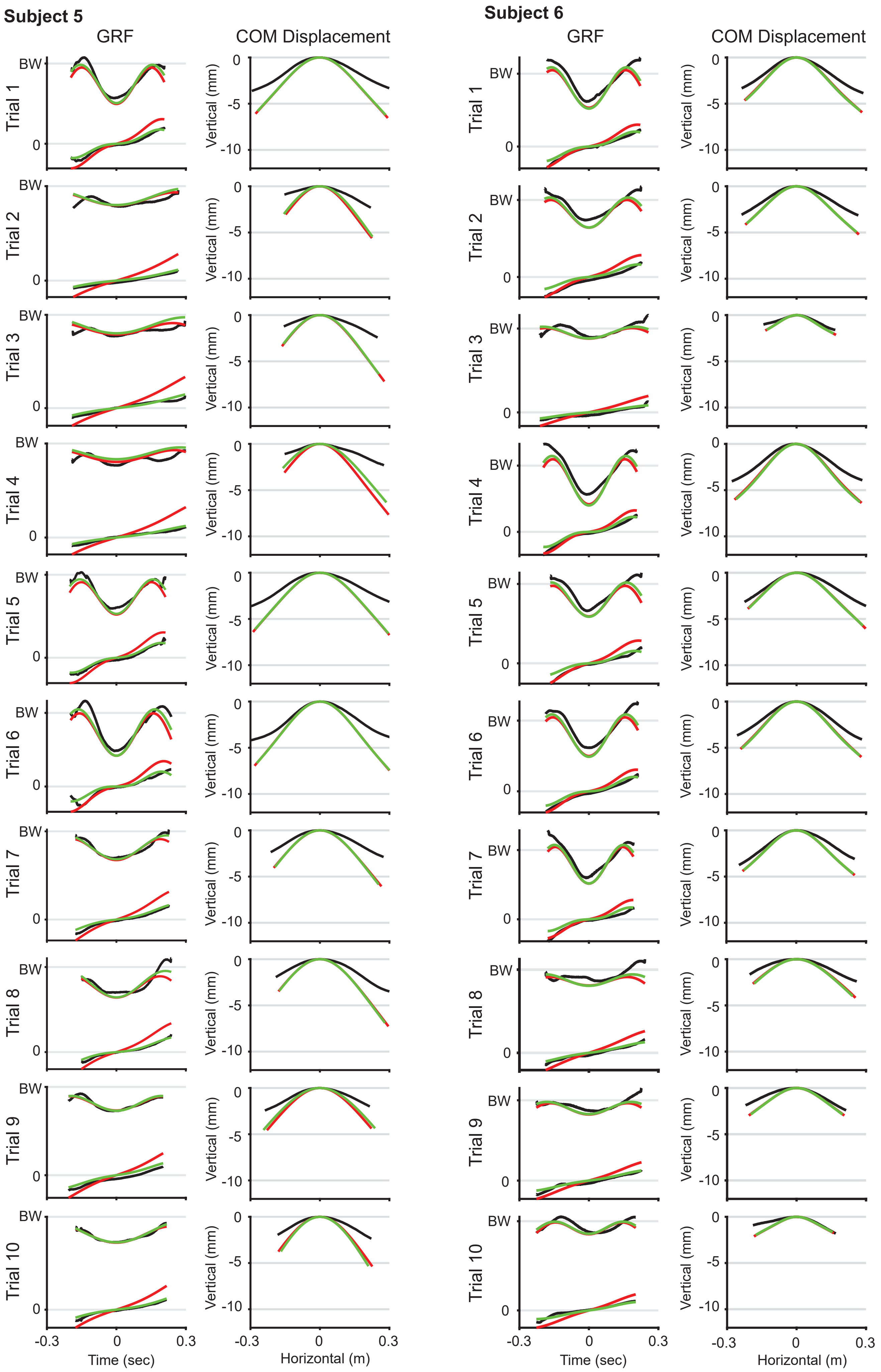


**Supplementary Figure 4: SLIP and ARSLIP fits to individual trials of Subjects 5 and 6.** For each panel, the experimental data is plotted in black, the SLIP fit is in red, and the ARSLIP is in green. For the GRF fits, the GRFs are normalized to subject weight and are plotted versus time. The COM displacement is plotted relative to the midstance position.

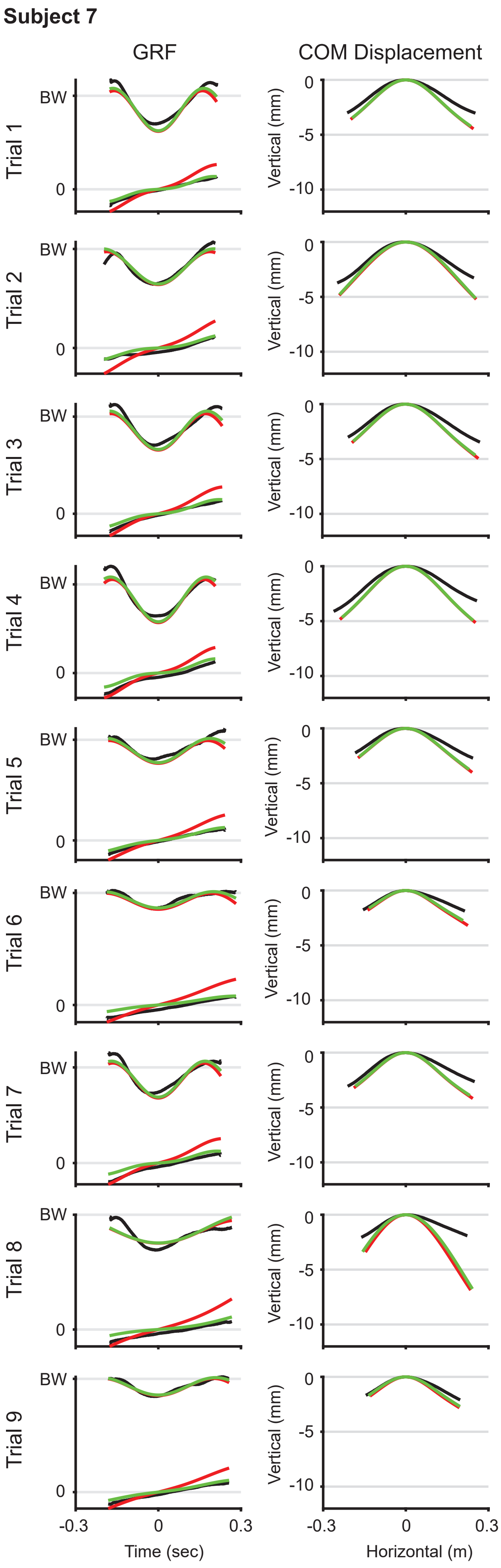


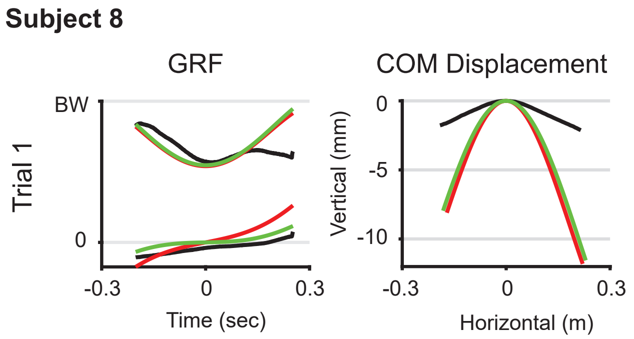


**Supplementary Figure 5: SLIP and ARSLIP fits to individual trials for Subjects 7 and 8.** For each panel, the experimental data is plotted in black, the SLIP fit is in red, and the ARSLIP is in green. For the GRF fits, the GRFs are normalized to subject weight and are plotted versus time. The COM displacement is plotted relative to the midstance position.
